# Linearised loop kinematics to study pathways between conformations

**DOI:** 10.1101/2021.04.11.439310

**Authors:** Antonius G.L. Hoevenaars, Ingemar André

## Abstract

Conformational changes are central to the function of many proteins. Characterization of these changes using molecular simulation requires methods to effectively sample pathways between protein conformational states. In this paper we present an iterative algorithm that samples conformational transitions in protein loops, referred to as the Jacobian-based Loop Transition (JaLT) algorithm. The method uses internal coordinates to minimise the sampling space, while Cartesian coordinates are used to maintain loop closure. Information from the two representations is combined to push sampling towards a desired target conformation. The innovation that enables the simultaneous use of Cartesian coordinates and internal coordinate is the linearisation of the inverse kinematics of a protein backbone. The algorithm uses the Rosetta all-atom energy function to steer sampling through low-energy regions and uses Rosetta’s side-chain energy minimiser to update side-chain conformations along the way. Because the JaLT algorithm combines a detailed energy function with a low-dimensional conformational space, it is positioned in between molecular dynamics (MD) and elastic network model (ENM) methods. As a proof of principle, we apply the JaLT algorithm to study the conformational transition between the open and occluded state in the MET20 loop of the Escherichia coli dihydrofolate reductase enzyme. Our results show that the algorithm generates semi-continuous pathways between the two states with realistic energy profiles. These pathways can be used to identify energy barriers along the transition. The effect of a single point mutation of the MET20 loop was also investigated and the predicted increase in energy barrier is consistent with the experimentally observed reduction in catalytic rate of the enzyme. Additionally, it is demonstrated how the JaLT algorithm can be used to identify dominant degrees of freedom during a transition. This can be valuable input for a more extensive characterization of the free energy pathway along a transition using molecular dynamics, which is often performed with a reduced set of degrees of freedom. This study has thereby provided the first examples of how linearisation of inverse kinematics can be applied to the analysis of proteins.

## 1 Introduction

One of the key problems in reducing the computational burden of protein simulation, structure prediction and design is effective sampling of the high dimensional conformational space [41]. What constitutes effective sampling depends on the goal. If the purpose is to find a minimum energy conformation, such as in *de novo* structure prediction, the full conformational space is within scope, while in other problems only conformations in the vicinity of a reference conformation are of interest, e.g. refinement [36] or analysis of conformational changes [39]. This paper addresses the latter type of sampling problems with a specific focus on loop motion analysis, which among other things enables the studies of conformational changes. Conformational changes are of central importance to the function of many proteins [17, 20, 16], including ligand binding [49] and enzymatic catalysis [31].

A direct way to improve sampling efficiency is to reduce the conformational search space by using an internal coordinate representation. An internal coordinate representation where only the backbone *ϕ* and *ψ* torsion angles are allowed to move reduces the conformational space without greatly compromising accuracy [18]. When a backbone is kinematically constrained, the conformational search space can be further reduced. For example, the conformation of a flexible loop is constrained by the start- and end residues, which are considered fixed with respect to each other. Loop closure thereby imposes six constraints: three translations and three rotations. Consideration of kinematic constraints will thus reduce the conformational search space by six.

The difficulty of considering loop constraints in an internal coordinate sampling algorithm is the transformation of the six loop closure constraints into the internal coordinate space [19, 34]. Finding values for the internal coordinates that satisfy the closure constraints is the problem of inverse kinematics for which various solutions exist. The Cyclic Coordinate Descent (CCD) method [9] is an iterative algorithm that repeatedly cycles through all torsion angles in a loop, solving in each step a one-dimensional projection of the kinematic closure problem in one torsion angle. The *KInematic Closure* (KIC) algorithm [33, 43], based on original work by Coutsias et al. [12], designates six torsion angles (three sets of *ϕ* and *ψ* torsion angles from three random residues) for loop closure. This separated closure problem has a finite number of solutions that can be solved analytically. Cortès et al. used a similar method that was developed by Renaud [42] to pioneer a rapidly-exploring random tree (RRT) method [11], which has subsequently led to a range of other algorithms [13, 6, 37].

However, while these algorithms have been game-changing, e.g. for loop modelling with fragments [27, 25, 48], they are not well suited to study conformational changes. That is because they do no permit close control of the sampling direction. In the case of CCD this is a result of the nested iterative scheme and the sequential approach of sampling and imposing constraints. In the case of KIC and RRT-based algorithms this is due to the artificial division of the conformational space into two subspaces; one dedicated to the inverse kinematics problem and one for exploratory sampling. For example, Al-Bluwi et al. discussed the need for additional filtering to ensure that subsequent samples are within a predefined distance from each other [2].

This paper complements existing solutions by introducing a novel method that enables tight control over constrained conformational sampling of a backbone segment. The main novelties of this paper are 1) a direct expression of a first-order approximation of the inverse kinematic relations of a protein backbone, i.e. its inverse Jacobian analysis, 2) a novel iterative loop closure procedure, and 3) a Jacobian-based Loop Transition (JaLT) algorithm to generate semi-continuous pathways between loop conformations. The presented inverse Jacobian analysis is performed with a method that was originally developed for the analysis of mechanisms with internal closed loops [24].The inverse Jacobian analysis presented here complements existing methods to analyse the standard (forward) Jacobian, which has been used in protein engineering to transform force vectors from Cartesian space to internal coordinates [38, 46], analyse the impact of changes to internal coordinates on the Cartesian coordinates [10] and analyse constrained motions [8].

The JaLT algorithm is positioned in between molecular dynamics (MD) and elastic network model (ENM) methods, which are often employed to study conformational transitions. MD-based methods are fully atomistic, set up to explore a large number of molecular degrees of freedom (DoFs), and rely on stochastic sampling. As a consequence MD-based methods are extremely time-consuming, even with advanced sampling techniques [26, 39]. ENM-based methods represent a protein as a network of residues and a simplified inter-residue energy function [26, 14, 8, 4]. As shown by Lu and Ma [32], additional intra-residue energy terms can be included, but energy models in ENM are still considerably less detailed than those utilized for MD. ENM-based methods are computationally efficient and have been very successful at predicting global conformational changes that involve large protein domains, but they are less suited to the studies of local changes such as loop conformational dynamics which involves subtle energy trade-offs [29]. The presented method aims to fill that gap with an algorithm that combines low-dimensional conformational sampling with the energetic detail of Rosetta’s all-atom energy function [3].

As a proof of principle, we apply the JaLT algorithm to study conformational changes in a loop from dihydrofolate reductase (DHFR). DHFR reduces dihydrofolic acid by transfer of a hydride from NADPH. The catalytic mechanism of the enzyme is tightly linked to the state of the MET20 loop near the active site of the protein [40]. Depending on the conformation of MET20 it can either open, close or occlude access of NADPH to the active site. Crystal structures corresponding to the three different states are available and show that the largest difference in conformation is between the open state and the occluded state, which is the transition we study in this paper. Mutational analyses have previously verified the importance of MET20 loop flexibility in the function of DHFR. For example, replacement of four residues in the loop with a single glycine resulted in a 500-fold reduction in hydride transfer [30]. We use MET20 as a model system to characterize the behaviour of the algorithm and generate example simulations for the full transition between the closed and occluded state in DHFR.

## 2 Results

The main result of this paper is the JaLT algorithm, which is introduced schematically in Fig. 1 and described in detail in the Method section. Linearisation of the backbone kinematics is central to the algorithm. This linearisation allows the transformation of vectors from Cartesian space to internal coordinates and vice versa. Thanks to this, all possible sampling moves can be expressed in the same reference frame which in turn enables a straightforward weighing of alternative moves. This is used to give greater weight to moves that are aligned with negative energy gradients. Finally, a linear model also enables a Newton-Raphson loop closure scheme, which ensures that loop closure is maintained throughout a simulation. As a result, the JaLT algorithm can simulate pathways that morph a start conformation into a target conformation without breaking closure constraints along the way.

**Figure 1:**
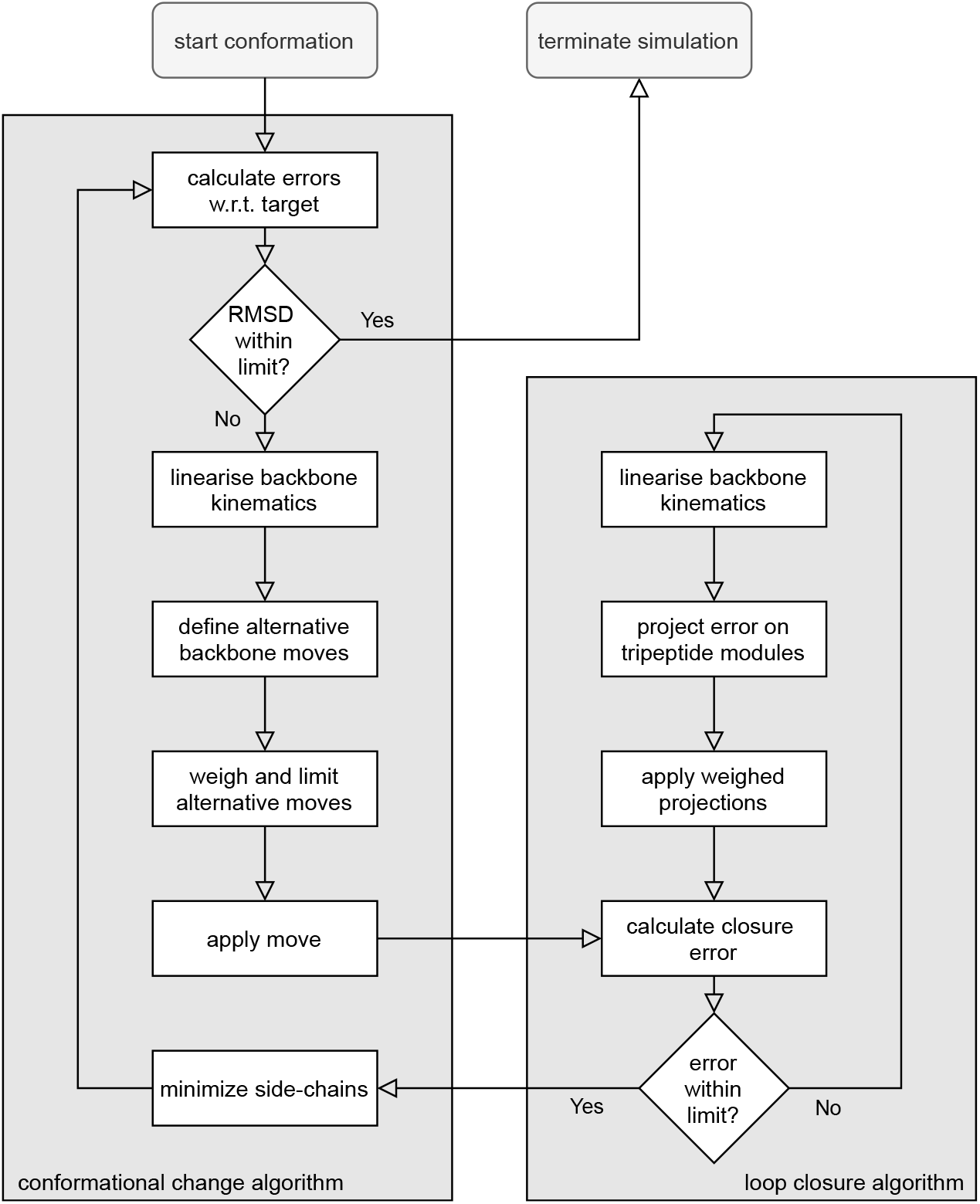
A flowchart giving a high-level overview of the Jacobian-based Loop Transition (JaLT) algorithm, which consists of a main conformational change simulation algorithm and a supporting loop closure algorithm, both based on a linearised model of the backbone kinematics.

The method shares the benefit of computational efficiency with ENM, since a JaLT simulation to generate a semi-continuous path from a start conformation to a target conformation of a loop requires just minutes on a single core of a 2.4 GHz Quad-Core Intel Core i5 processor. On the other hand, it enables exploration of energy landscapes in greater detail through the integration of an all-atom energy function, which brings it closer to MD simulations.

In this Results section, first the characteristics of the developed algorithms are described, followed by a presentation of simulation results that illustrate the validity of the method. Next, the use of the JaLT algorithm is demonstrated by an application to the MET20 loop in DHFR. We demonstrate how the JaLT algorithm can be used to find minimal energy pathways between conformational states and how it can help to identify torsion angles that are critical for the conformational change (often referred to as collective variables).

### Linear analysis methods

The developed algorithms are built around a linearisation of the backbone kinematics. Tripeptides are used to organize the linearisation. A tripeptide contains six dihedral angles, which thereby enables a one-to-one mapping between Cartesian coordinates and internal coordinates (see Method section for more details). The N-atoms that connect subsequent tripeptide segments are marked as *key atoms*.

Linearising a system comes with advantages and a disadvantage. The disadvantage is that a linearised model is only valid in the direct vicinity of the instantaneous conformation. As such, a simulation must always start from a valid, kinematically closed start conformation. To sample within the region where the linear approximation is valid, the JaLT algorithm scales a move based on a (configurable) maximum change in torsion angle per move, which is further scaled down by the algorithm in case an invalid move is detected. The advantages are:

- *Samples form a semi-continuous path.* The root-mean-square deviation (RMSD) of CA atoms of subsequently sampled conformations is within few hundreds of Å so that they are directly connected in the conformational space. As a result, subsequently sampled conformations together form a semi-continuous pathway from the start conformation to the target conformation.
- *Ability to consider closure constraints while defining a move*. This avoids exploration of moves that are subsequently rejected at the scoring phase. Because scoring a conformation is computationally costly, avoiding unproductive moves is a big computational advantage.
- *Great freedom to distribute the torsional changes in each move*. The redundant DoFs in a loop can be used to optimize a move based on specific criteria. The JaLT algorithm uses this freedom to assign larger weight to segments of the loop where the desired changes aligns with a negative energy gradient. The fact that linear algebra can be used makes it straight-forward to perform weighing.
- *Deterministic analysis method to generate semi-continuous pathways*. Because the method does not rely on stochastic sampling, a simulation can be fully deterministic. This enables the design of controlled experiments. In the JaLT algorithm a stochastic factor is introduced to sample alternative pathways, but this is not intrinsic to the modelling method and can also be optimised or customised for other purposes.

#### Algorithm objectives and options

The linearisation of the conformational space enables detailed control of the direction of moves while sampling in a deterministic manner. In this study we use this flexibility to achieve two different objectives:

1. **Converging on target.** The definition of moves in the main loop of the JaLT algorithm (see Algorithm 1 in the Method section) combines kinematic and energy gradient information in a way that pushes a closed loop towards a target conformation along negative energy gradients where possible.
2. **Maintain loop closure.** A move that is based on a first-order approximation of the kinematics introduces small closure errors, which will quickly result in runaway behaviour if not controlled. Therefore, each iteration of the JaLT algorithm ends with a call to a novel loop closure algorithm. This iterative loop closure algorithm is built around the same linearised kinematic model and brings the kinematic loop closure error down to within specified margins. This loop closure step thus acts as a kinematic feedback loop on the conformational changes defined by the first objective.

There are several user-defined options that can be used to control the behavior of JaLT, shown in Table 1. The impact of these options are outlined below and in Algorithm 1 in the Method section.

**Table 1:** User options to control the JaLT algorithm.

| option | description | default value [unit] |
| --- | --- | --- |
| it_max | number of iterations after which algorithm is stopped if not converged | 5000 [-] |
| pdb_interval | interval at which PDBs are stored | 10 [-] |
| hor_max | maximum number of tripeptide segments at either side of a key atom that are involved in a key atom move | 3 [-] |
| rmsd_lim | RMSD of CA atoms with respect to the target pose at which the algorithm is stopped | 1.0 [Å] |
| torsion_step_max | maximum torsion change during a single move | 10 [deg] |
| w_scaling | scaling factor for weighing energy gradient of alternative moves | 0.1 [-] |
| w_torsion | relative weight given to torsion errors instead of Cartesian error | 0.5 [-] |

### Algorithm performance

This section benchmarks the JaLT algorithm and illustrates the impact of various parameters in the algorithm. The presented analyses are performed on residues 9-24 in the MET20 loop of DHFR (PDB entries 1ra2 and 1rx7). The raw data is available in Supplemental Data 1.

#### Validation of linearisation

To validate the linearisation method that is used to define moves, firstly it is verified that the maximum torsion change of a move can be controlled. For different settings of the nominal step size (using `torsion_step_max`) Fig. 2a shows distribution graphs of the maximum torsion angle change after each finished move (i.e. including a call to the closure algorithm). For each step size setting, ten simulations were performed, each consisting of 1000 loop iterations. The results are also summarized in Table 2.

**Figure 2:**
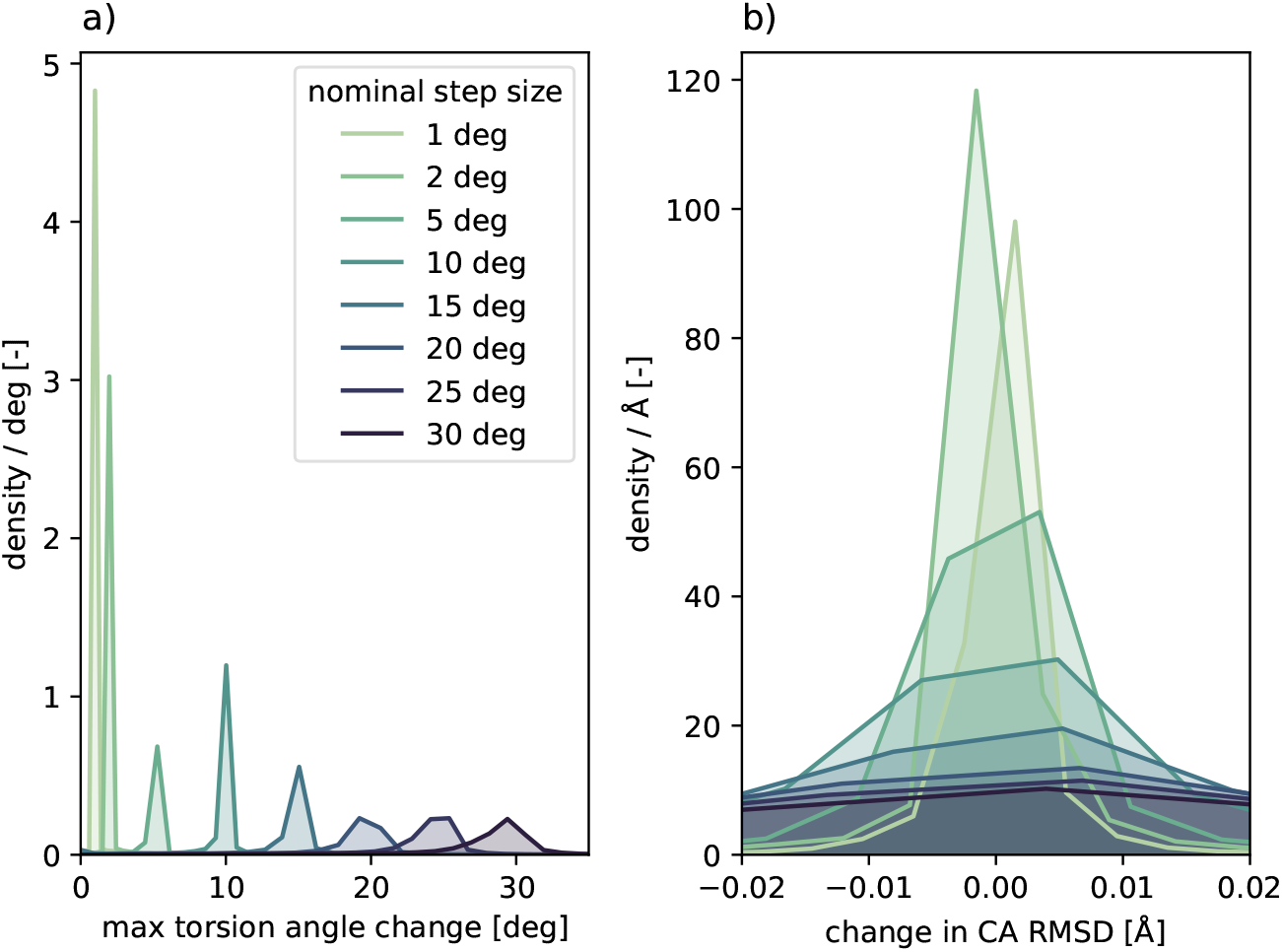
Density plots of the maximum change in torsion angle and the change in CA RMSD after each move, for desired maximum step sizes between 1 and 30 degrees.

**Table 2:** The mean error and the standard deviation (STD) of the maximum torsion angle change Δ*q_max_* per sampling step show that the algorithm controls the change in torsion angle with an approximate error of 10%.

| torsion_step_max [deg] | mean($\Delta q_{max}$) [deg] | inaccuracy of mean [%] | STD($\Delta q_{max}$) [deg] |
| --- | --- | --- | --- |
| 1.0 | 1.11 | 11.1 | 0.73 |
| 2.0 | 2.18 | 9.1 | 1.13 |
| 5.0 | 5.22 | 4.3 | 2.09 |
| 10.0 | 9.93 | -6.5 | 2.53 |
| 15.0 | 13.88 | -7.4 | 4.62 |
| 20.0 | 18.94 | -5.3 | 4.15 |
| 25.0 | 23.23 | -7.1 | 5.11 |
| 30.0 | 27.53 | -8.2 | 5.83 |

It can be observed in Table 2 that the mean of the maximum torsion angle change is within approximately 10% from the user-defined nominal step size. This close correlation between `torsion_step_max` and the actual mean error is also visualised in 3a. Moreover, the standard deviation of observed maximum torsion angle change also varies approximately linearly with the user-defined setting. This is shown in Fig. 3b, which reflects the sharper density peaks in Fig. 2a for smaller nominal step size settings. Up to a nominal step size of 10 degrees the mean of the actual maximum change is within one degree of the set value and the standard deviation within three degrees.

**Figure 3:**
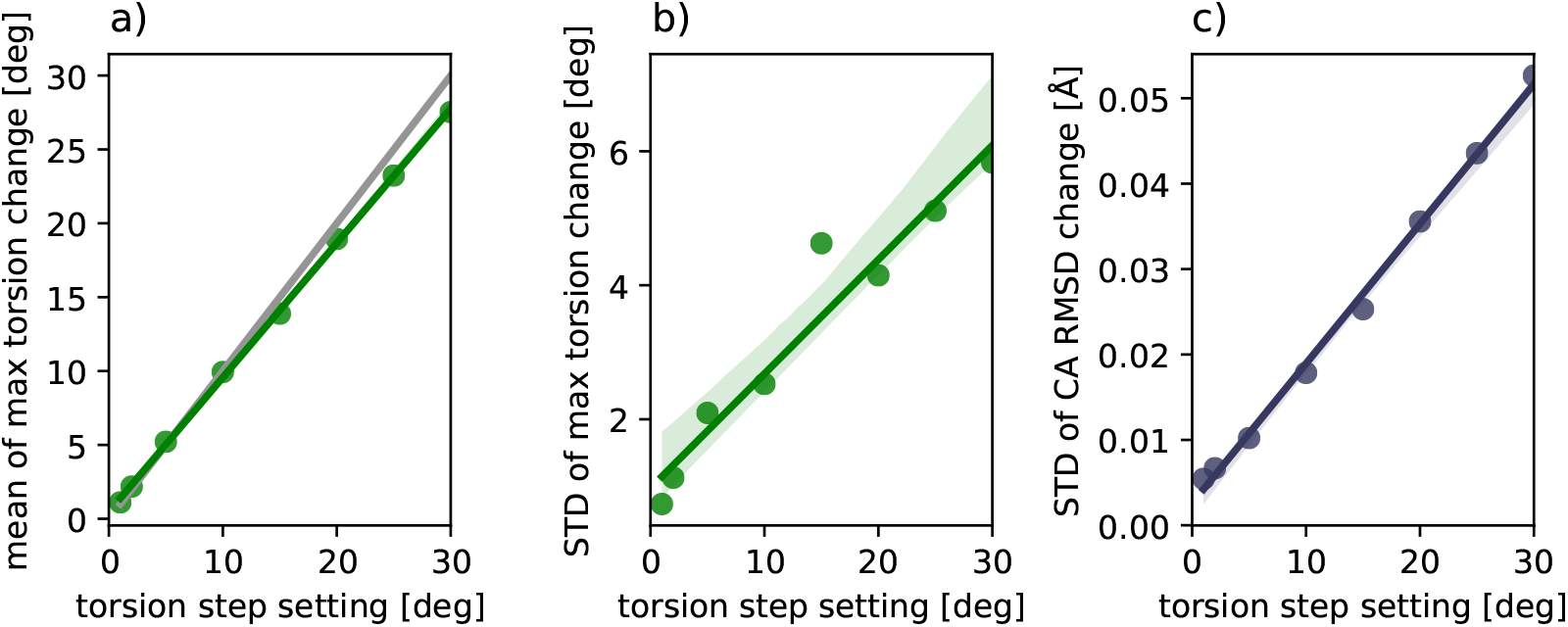
Correlation between user-defined torsion step size setting and actual maximum torsion change after each move followed by loop closure, as well as a correlation with the standard deviation (STD) of actual maximum torsion changes and STD of changes in CA RMSD. The grey line in a) represents a perfect correlation.

However, control over the torsion angle change is not the goal on itself. The goal is to control the conformational change, which is better reflected in the RMSD of the CA atoms of the loop. Figure 2b and Fig. 3c show that changes in CA RMSD for each step are within a few hundredths of an Ångström. Conformations with a CA RMSD distance between them in the order of hundredths of an Ångström are considered to form a semi-continuous pathway. This therefore validates that controlling the maximum torsion angle changes between samples in the JaLT algorithm to within tens of degrees enable sampling from a semi-continuous pathway.

#### Convergence of algorithm

To study how the JaLT algorithm converges on a target conformation, we analysed trajectories consisting of 1000 iterations, with the CA RMSD of the target conformation 3.8 Å away from the starting conformation. Figure 4 shows the distribution of the minimum RMSD values encountered during each simulation, as a function of step size. Again, each simulation was repeated ten times. RMSD values are calculated for respectively all torsion angles and all CA atom positions in comparison to the target state. Moves were either defined based on errors with respect to the target state in Cartesian coordinates of the key atoms (`w_torsion=` 0.0) or in torsional values of the backbone dihedral angles (`w_torsion=` 1.0).

**Figure 4:**
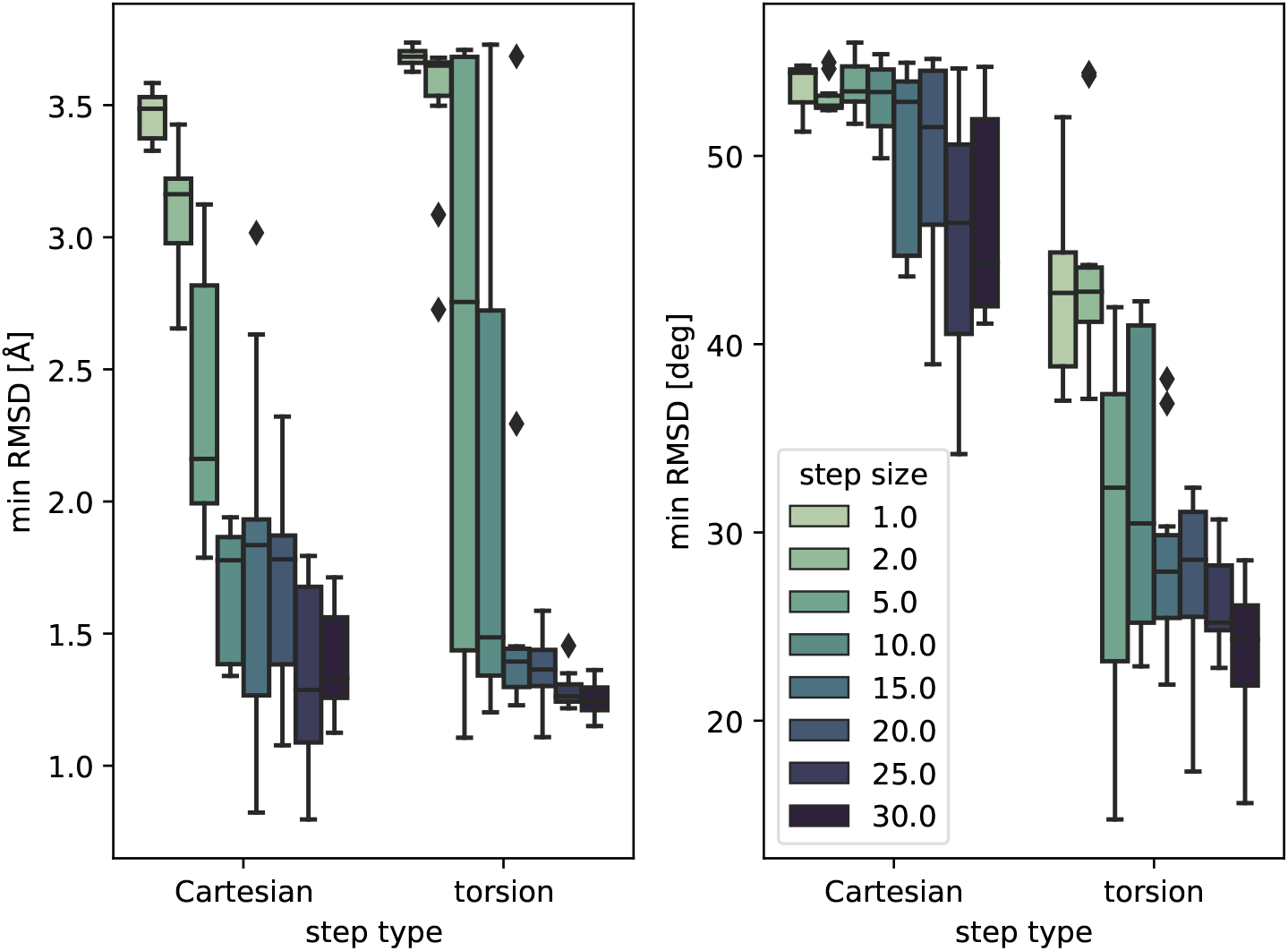
Box plots of the minimum RMSD for the CA atoms and for the torsion angles with respect to the target reference structure during simulations of 1,000 iterations for `w_torsion` either set to 0.0 (cartesian) or to 1.0 (torsion).

Figure 4 shows that for step sizes smaller than 5 degrees the algorithm does not manage to move very far from the start conformation within 1000 iterations. Step sizes of 10 degrees or more are required for significant progress. It can also be observed that moves based on Cartesian error have a relatively small effect on the torsion error, but moves based on torsion error have an effect on the CA atom RMSD that is comparable to moves based on Cartesian error. This is further investigated later in this paper.

#### Effect of using energy gradients when selecting moves

Figure 5 presents an analysis on the effect of different forms of weighing of alternative moves. The weighing in the JaLT algorithm is implemented based on the energy gradient of the Rosetta energy function, and to assess its effectiveness it is compared to simulations where weighing is based on kinematic errors alone (which is how weighing is implemented in the loop closure part of the algorithm). Simulations were carried out with a step size of 10 degrees and were continued for 5,000 iterations. Figure 5 shows that weighing based on energy gradient significantly reduces the energies of the sampled conformations, without negative impact on the RMSD of the resulting conformations.

**Figure 5:**
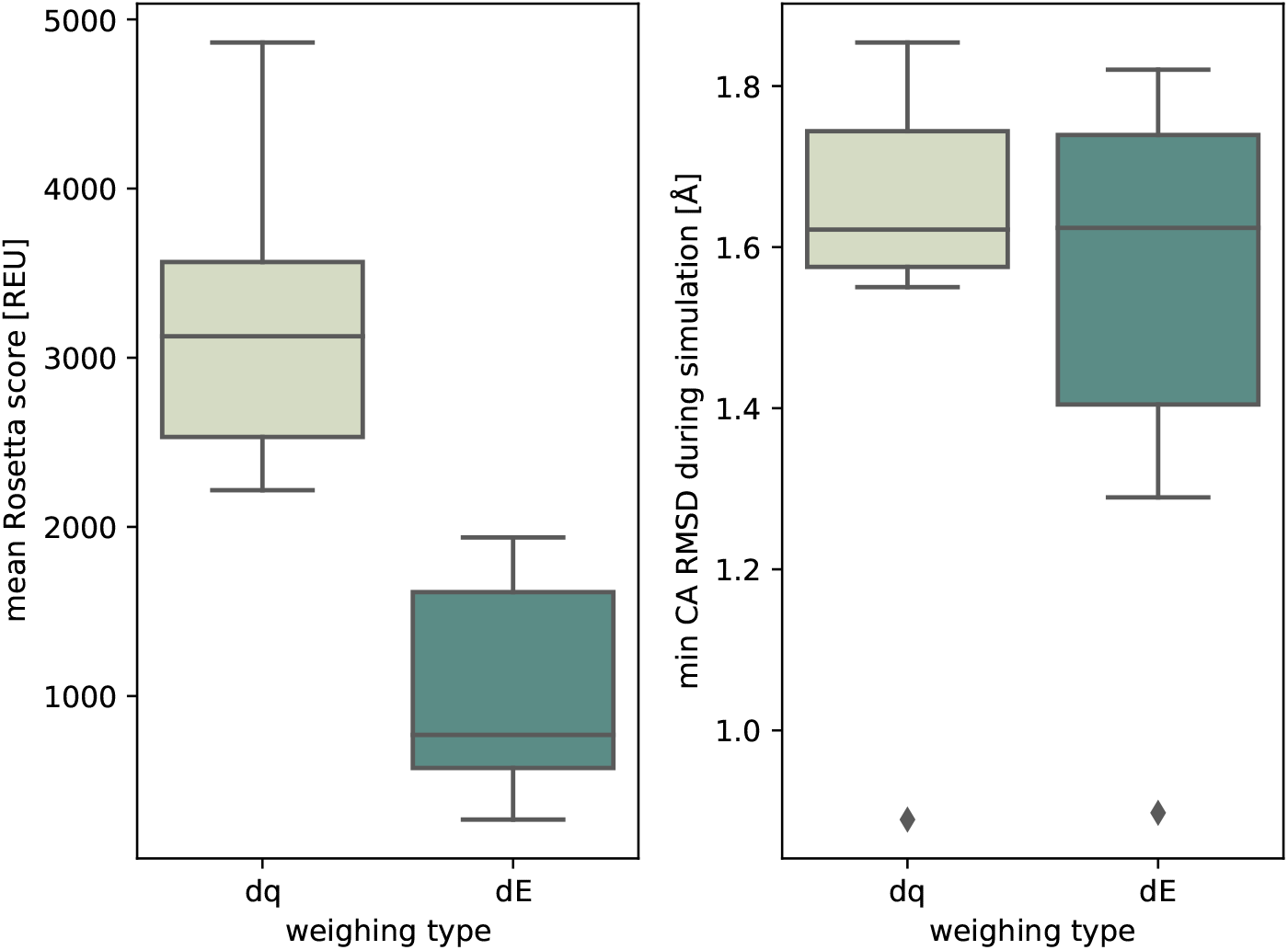
Linearisation enables weighing of alternatives moves, for example based on the energy gradient of the move (dE) or the inverse of the required torsion angle (dq) change which significantly improves the mean energies of sampled pathways without negatively affecting the CA RMSD.

#### Optimal combination of Cartesian and internal coordinates in loop transition analysis

In this section we investigate how Cartesian and torsion information can be combined to improve loop transition simulations. This can be controlled via the option `w_torsion`, which is the relative weight given to torsion errors over Cartesian errors in the definition of moves. Figure 6 shows the result of JaLT simulations where the impact of `w_torsion` on the CA RMSD and the torsion angle RMSD is studied. The analysis demonstrates that dedicating part of the move to reducing the torsion angle error initially reduces the lowest sampled RMSD, but it increases again as the torsion error weight factor approaches 1.0. A similar, but less pronounced effect can be seen for the torsion angle RMSD. Combination of kinematic information can thus increase performance in terms of convergence to a target conformation.

**Figure 6:**
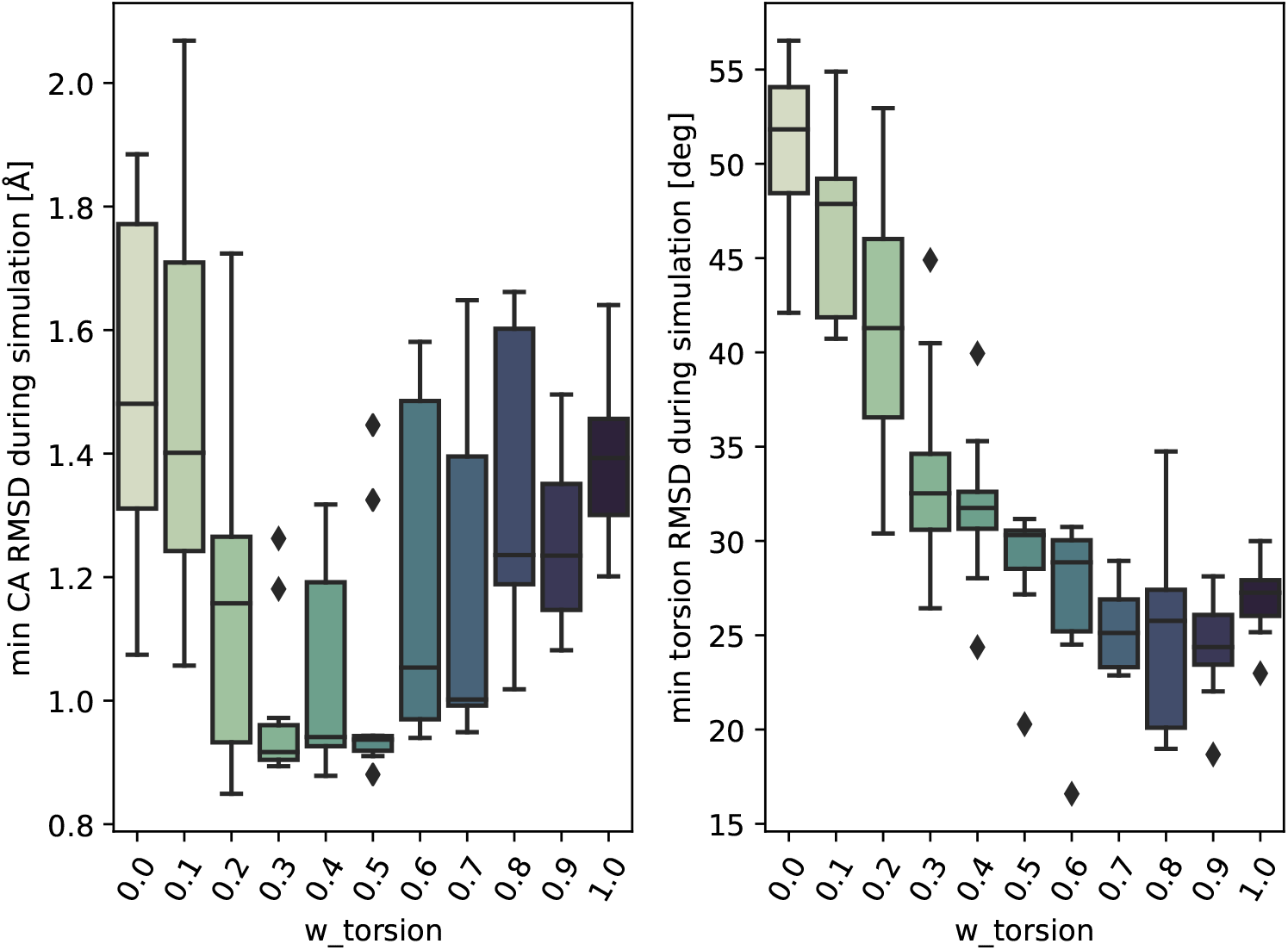
A full Jacobian analysis enables a simultaneous consideration of errors defined in Cartesian space and torsion space and thereby enable the optimal use of all available information.

While construction of kinematically accessible pathways is an important objective, we also want to constrain sampling to transitions that are energetically feasible. Figure 7 shows the effect of `w_torsion` on the mean energy during a simulation. A minimum can be observed around `w_torsion=`0.5 which strengthens the observation that a combination of information from Cartesian coordinates and torsion values improves performance. Figure 8 explores the combined space of energy, CA RMSD and weighing (`w_torsion`) found in Figure 6 and Figure 7, confirming that giving equal weight to Cartesian errors and torsion angle errors is approximately optimal.

**Figure 7:**
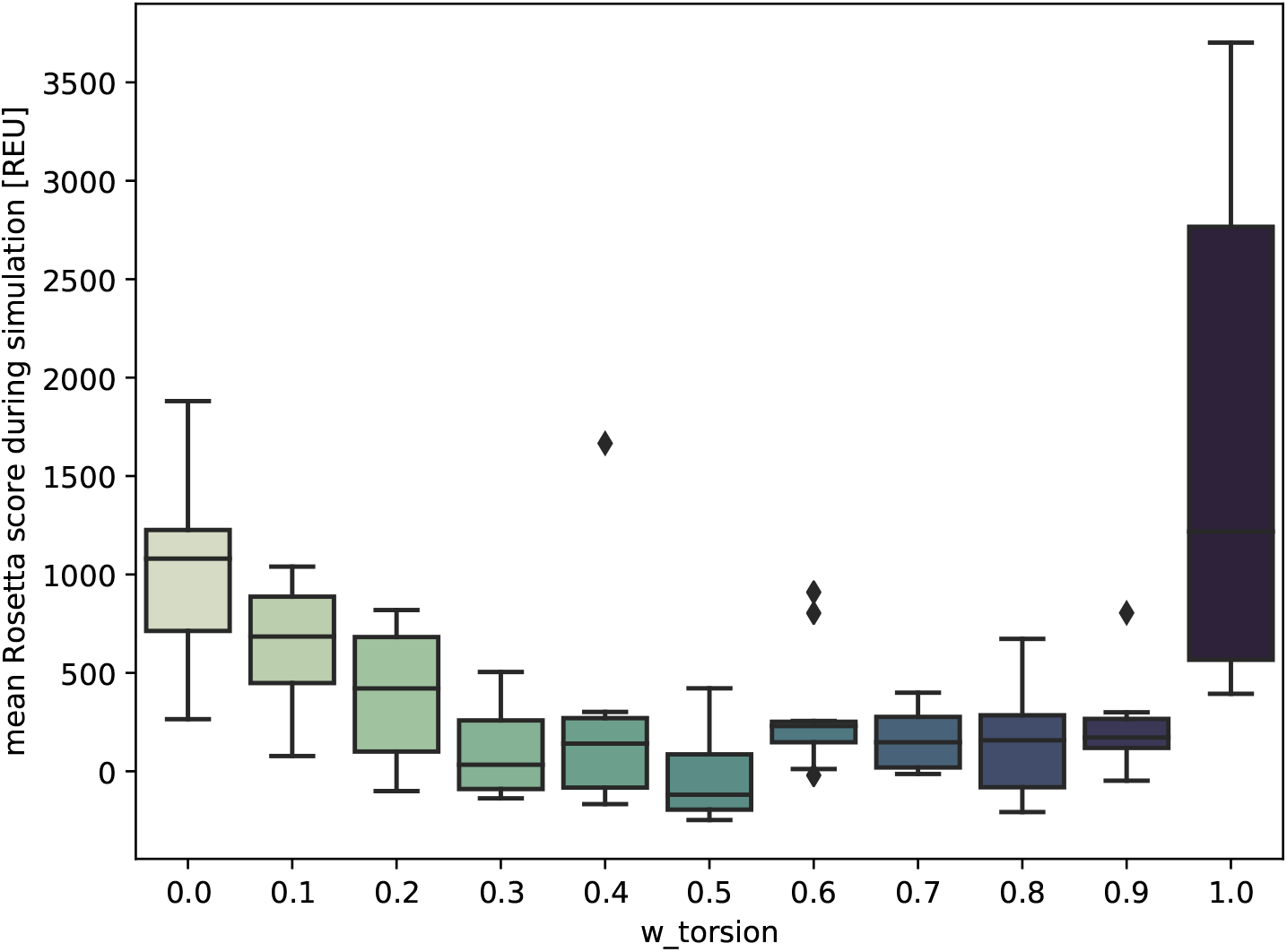
Combining Cartesian information and torsion information enables the algorithm to find energetically more favourable pathways.

**Figure 8:**
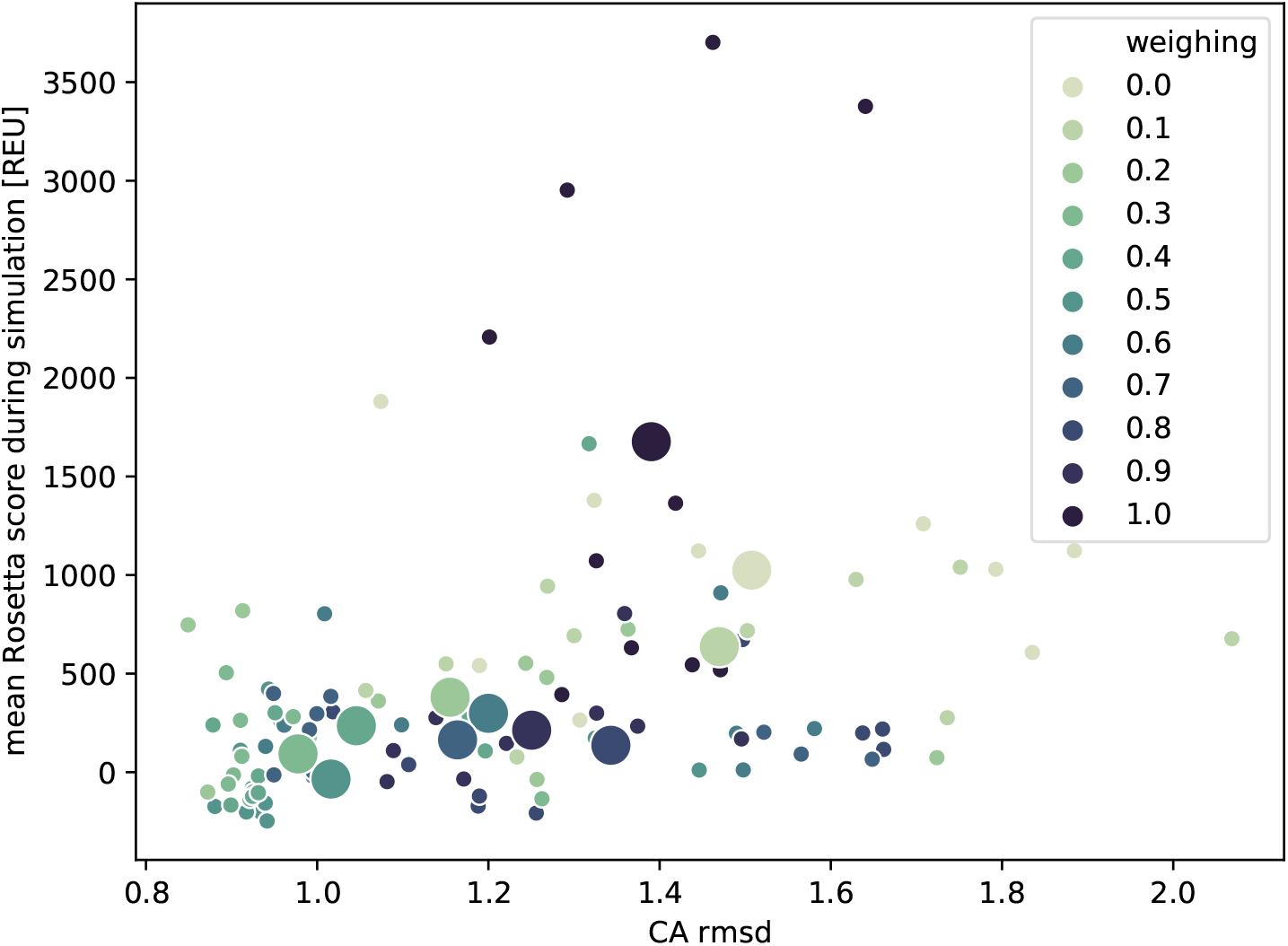
A scatter plot of the RMSD for the CA positions versus the mean score of simulations, with the mean values for each different weighing added with a large circle. For the presented conformational change analysis the optimal mixture of Cartesian and torsion angle information is approximately 50-50.

### Application of the JaLT algorithm to modelling of loop transitions in DHFR

In the previous section it was shown that the JaLT algorithm can sample conformations from semi-continuous pathways by controlling the maximum torsion angle change between conformations. It was also shown that energy gradients can be used to reduce the energy of a move, while combining Cartesian and torsion information improves convergence to the target state.

In this section we demonstrate how the methodology can be used to investigate conformational changes of loops in proteins. Three examples are given for possible applications of the JaLT algorithm: exploration of pathways between conformational states, identification of collective variables, and analysis of the effect of mutations on the conformational landscape. All results are generated using the MET20 loop of DHFR.

To characterize pathways between the open and occluded state of MET20 we ran JaLT simulations starting from the open state (PDB 1ra2) with the occluded state (PDB 1rx7) as the target. The simulations were run with equal weight on Cartesian and torsion value errors (`w_torsion=`0.5) and ran until the instantaneous conformation was less than 1 Å CA RMSD from the target. For this paper only simulations that converged within 500 iterations were accepted, because shorter pathways are more direct which makes the results easier to visualise. The raw data can be found in Supplemental Data 2.

#### Exploration of pathways between conformational states

Figure 9 shows the trajectory with the lowest energy barrier from a set of 2000 started simulations. Around iteration step 150 of the example simulation, the transition goes through a high energy barrier. Once this barrier is traversed the RMSD drops rapidly. The barrier is associated with steric clashes between the sidechain of Trp22 and the backbone of Ile14 and Gly15. The carbonyls of Ile14 and Gly15 are also momentarily clashing while steric repulsion is observed between Gly15 and Glu17 (and to a lesser extent Met20). The release of energy from the top of the barrier is mostly associated with a 35 degree change in the *ϕ* value of Gly15, which reduces steric repulsion. The complete series of PDB files generated at intervals of 10 iterations for the presented simulation can be found in Supplemental Data 3.

**Figure 9:**
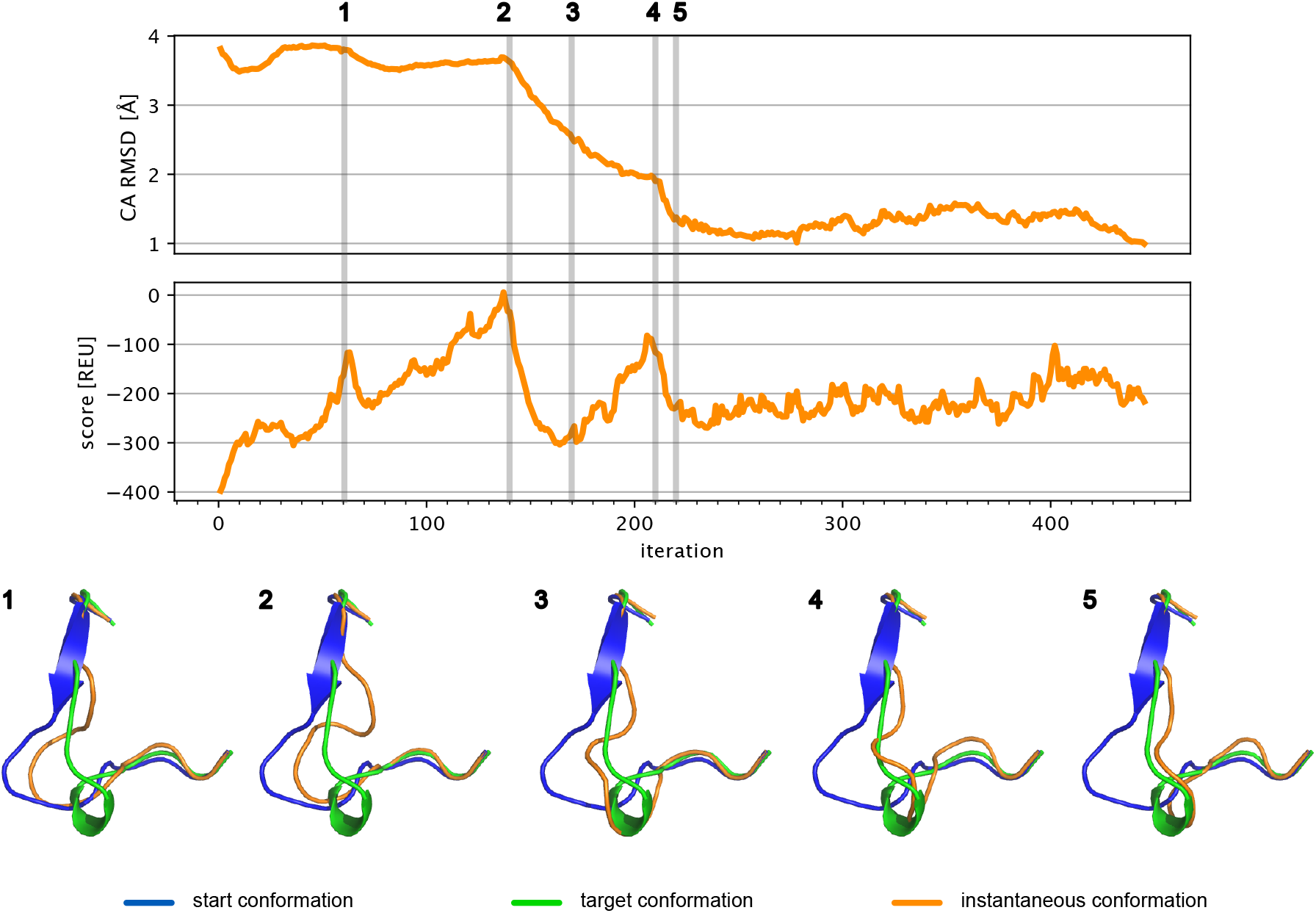
The trajectory with the lowest maximum Rosetta Energy Unit (REU) score that was generated for the loop transition of the MET20 loop from the open state (PDB 1ra2) to within 1 Å of the closed state (PDB 1rx7) together with snapshots of the loop structure at various stages of the simulation.

#### Identification of collective variables

An analysis of the torsion angle values could help to identify the dominant torsion angles for the change between two loop conformations. Such analysis is common as a preparatory step for biased MD simulations. Figure 10 shows an analysis of the torsion angles that resulted from the simulation introduced in Fig. 9. The results indicate clear differences in the variation of torsion values at different positions in the loop during the simulation. Wu & Post [50] studied the MET20 conformational transition using their MD-based adaptively biased path optimization method and found that *ψ*14, *ψ*18, and *ψ*19 were good choices for collective variables based on the observation that their values differ the most between start and target conformation. However, some torsion angles that have comparable values in the start and target conformation may need to adopt diverging values during the transition. Our simulation suggests that torsion angles may go through a full 360 degree rotation during the transition (e.g. *ϕ*15) or a temporary rotation (e.g. *ϕ*17). A JaLT analysis smay thus be used as an initial approach to identify meaningful collective variables for more extensive and computationally intense MD simulations.

**Figure 10:**
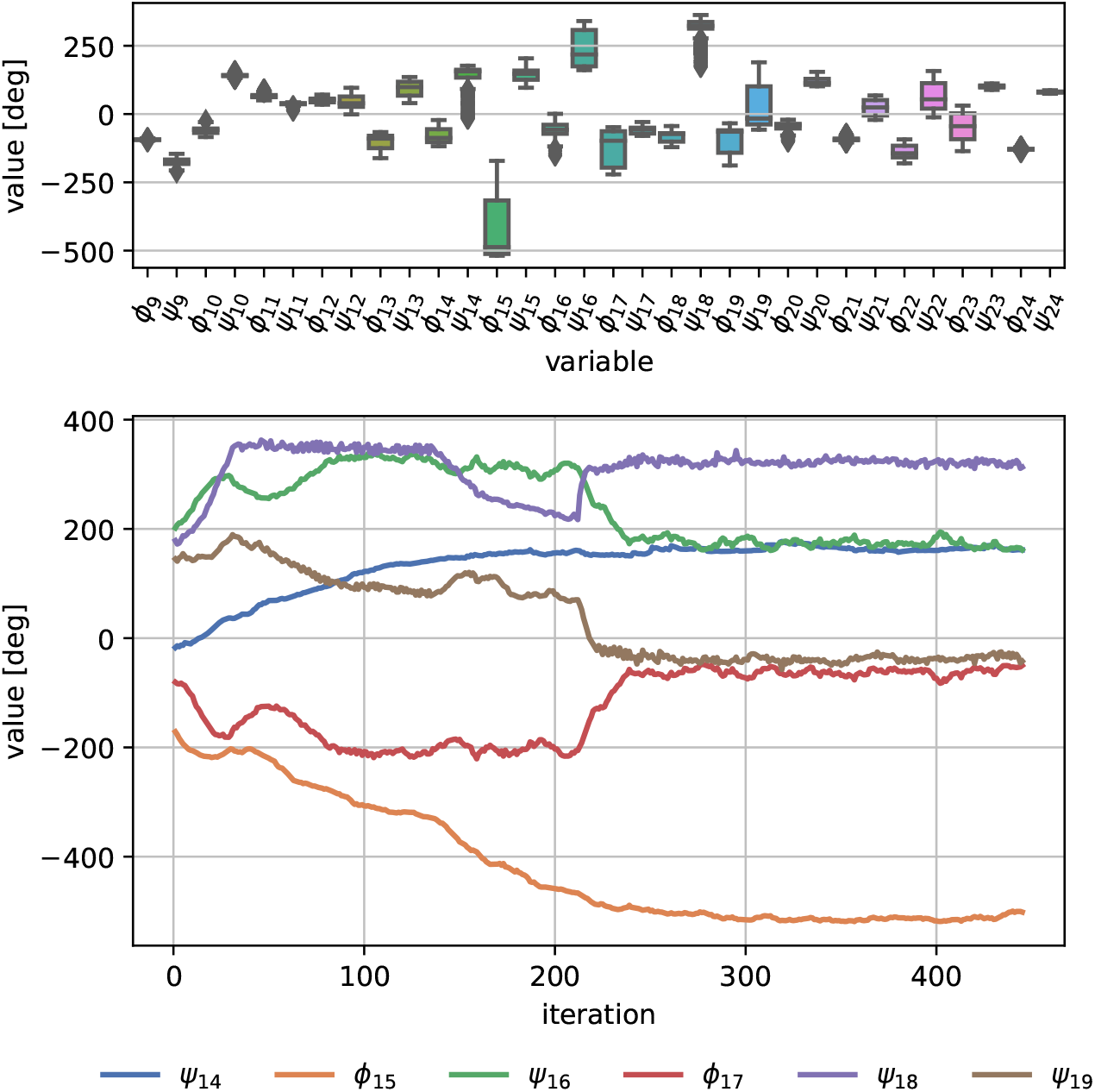
An analysis of the torsion angles throughout a simulated trajectory can identify possible collective variables. Interestingly, some torsion angles end up with approximately the same value as they started with, but go through significant temporary changes during the transition.

#### Analysis of conformational change landscape

Kinetic data on DHFR have demonstrated that mutations in the MET20 loop can have significant impact on the activity of the enzyme. Mutations of I14 to Val, Ala and Gly results in significant reduction of hydride transfer rate [44], with the Gly mutation having the biggest impact. Quantum mechanics-molecular mechanics simulations suggest that mutations of I14 leads to increased free energy barrier for hydride transfer [15]. To analyse whether JaLT could be used to investigate this phenomenon, loop transition simulations were also performed with a I14G variant of DHFR, and the result is compared with the wild-type protein.

In Fig. 11 the simulation with the lowest energy barrier is shown for both the wild-type and the glycine-mutated variant. The progress of the simulated trajectories is expressed using Δ*D_rmsd_* as described by Arora and Brooks [5]. The expression for Δ*D_rmsd_* is repeated here in adapted form for convenience:

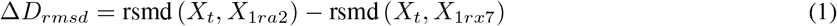

**Figure 11:**
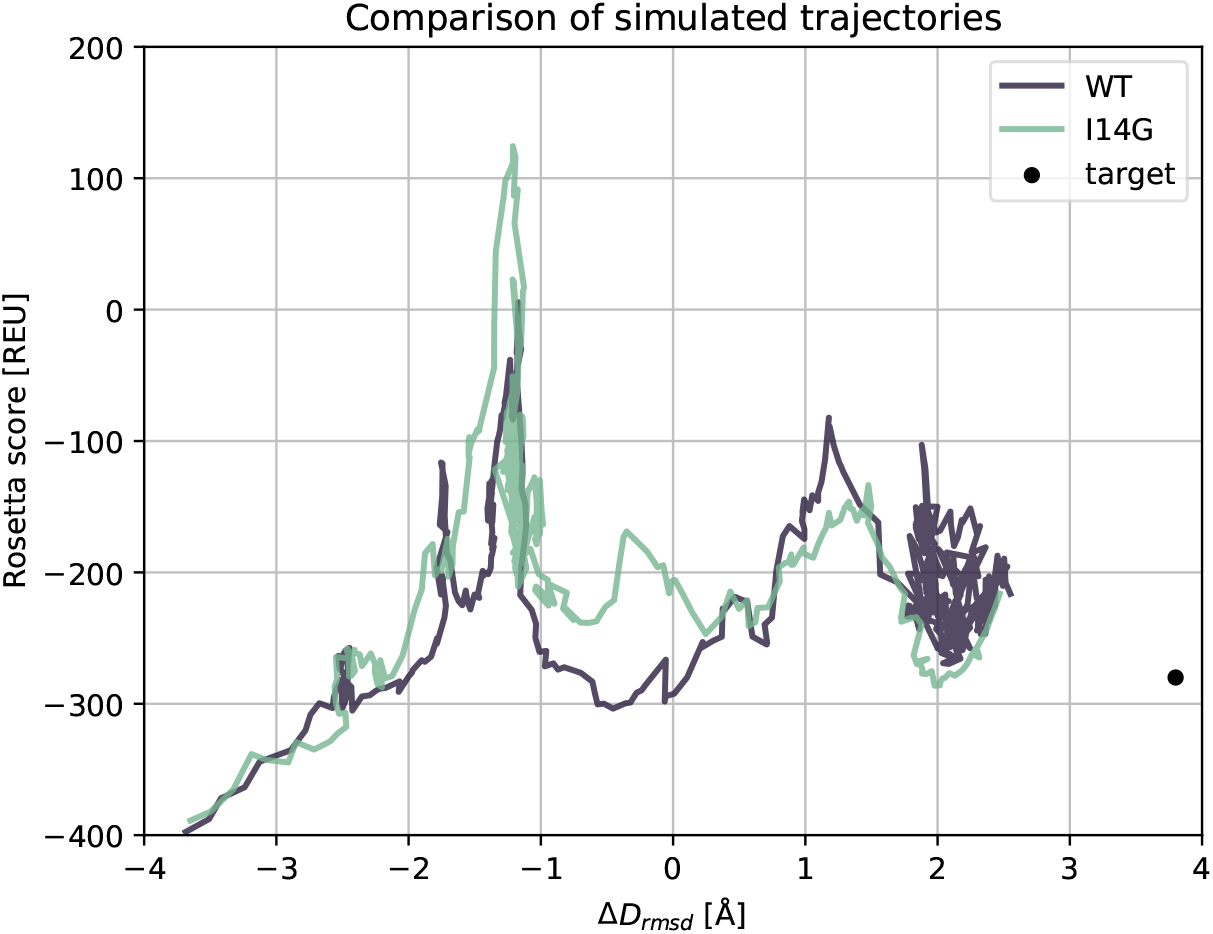
Conformational change is guided by energy, and simulations of a conformational change of the MET20 loop in its wildtype and I14G mutated form show clear differences in the region midway between the two conformations. Note that because the simulation is also driven by the gradients in torsions and energy, changes in RMSD does not have to be monotonic.

where *X_t_* is the instantaneous structure during simulation, *X*_1*ra*2_ and *X*_1*rx*7_ are the reference open and occluded states of DHFR, respectively.

The introduction of I14G leads to a higher energy barrier between the open and occluded state, which is consistent with the lower catalytic rate for this mutant. The barrier is associated with a clash between the carbonyl of Arg12 and the backbone of His124 on an adjacent loop. The release of energy at the top of the barrier is mostly associated with changes in the *ϕ*-angles of Ala9 and Val10. The complete series of PDB files generated at intervals of 10 iterations for the presented simulations can be found in Supplemental Data 3.

## 3 Discussion

This paper set out to demonstrate that linearisation of protein backbone kinematics is a valuable tool for the analysis of proteins. Figures 2 and 3 showed that for the example of the MET20 loop of DHFR the maximum torsion change of a closed loop move can be controlled with a standard deviation of a few degrees. This indicates that the loop closure algorithm only needs to make small corrections to the torsion angles that were calculated using the Jacobian transformations. This conclusion is supported by the observed changes in CA RMSD, which increase approximately linearly with the user-defined step size. Furthermore, the changes are small with deviations of only 0.04 Å even for a step size of 30 degrees. These findings are a first demonstration that a linear model of a protein backbone expressed in a low-dimensional set of internal coordinates can be used to sample conformations along a semi-continuous pathway.

While a smaller step size allows tighter control over the sampling, it can prohibit conformational exploration. With small step sizes (below 5 degrees for the example analysis in this paper) JaLT only explores a very local environment and does not progress towards the target, which is seen in the high RMSD values for CA and torsions in Fig. 4. A step size of 10 degrees results in a full transition towards the target, while being small enough to provide accurate linearisation and sample on a semi-continuous pathway.

One of the benefits of a linearised model is that vector algebra can be used to weigh alternative moves. Figure 5 showed that a weighing of alternative moves based on the energy gradient is effective in promoting lower energy moves. Because the value for the energy gradient is unbound, the values were mapped onto the range between zero and one. In this study we used a sigmoid function for this conversion, where the parameter `w_scaling` was introduced to give the user control over the shape of the sigmoid. Another example of weighing is the combination of information stored in Cartesian and internal coordinates, for which the parameter `w_torsion` was introduced. It was shown that a factor of 0.5 is roughly optimal for the given example. Possibly this is because moves based only on Cartesian errors tend to get stuck in confined regions of the conformational space, which can be overcome by also considering errors in dihedral angles. On the other hand, using only internal coordinate information will not lead to valid results because the anchors of the loop in the start- and target conformations are always slightly different, and these slight differences can have big impacts on dihedral angles due to the non-linear nature of the loop kinematics.

The JaLT algorithm was used to study the transition between the open and occluded state of the MET20 loop in DHFR. The algorithm is iterative and at each instantaneous conformation it favours moves that are beneficial from a perspective of energy, while moving the conformation towards the target. It often happens that the algorithm ventures into a direction that leads to a region of clashes and high energies, at which point the algorithm will not back up, but instead work its way through the region of high energy. We built a stochastic term into the model, which means that alternative pathways are explored in each trajectory. As such, by running more simulations the energy landscape of the loop transition can be explored in order to identify pathways with low barriers and to identify dominant pathways. An analysis of the flow between start and target state could then be used to characterize the transition. Alternatively, structural samples along the transition can be used as starting points for more extensive MD simulations to more accurately investigate free energy barriers of loop transitions.

The simulated pathway shown in Fig. 9 does not monotonically decrease the RMSD to the target conformation, i.e. at certain points along the trajectory it moves away from the target conformation before coming closer again. This behaviour is not surprising because the constrained kinematics of loops are highly non-linear. In order to comply with loop constraints and avoid clashes, it is easy to imagine that the RMSD of a loop may temporarily need to increase to move parts of that loop around a kinematically inaccessible or energetically unfavourable region.

This paper used the linearised model of the protein backbone kinematics for the analysis of conformational change, but the opportunities are broader. In the field of robotics, the linear algebra that is at the heart of this paper is extensively used for dynamic analyses such as stiffness analysis and natural frequency analysis [7, 47, 45, 23, 21] and has also been applied for the analysis of mechanisms with more complex, internally connected, kinematic networks [28, 22]. Such internally connected mechanisms are not unlike protein segments that are interconnected via hydrogen bonds, as for example investigated by Budday et al. [8], and may proof a new source of inspiration.

This study has focused on the methodological basis for loop transition analysis, but there are many ways in which the current methodology can be extended. The stochastic element of the simulations could be replaced by an energy-based Monte Carlo sampling with a Bolzmann criteria. Additional stochastic sampling of additional degrees of freedom can be introduced, such as *ω* and bond angles. Each step in the algorithm involves a round of minimization of side-chain torsion values, but more extensive optimisation can be added if desired. Also, there are many alternative ways for weighing of moves to be explored.

Finally, the methodology is not confined to studies of conformational changes in loops. Inverse Jacobian analysis is perfectly positioned to model kinematically constrained systems like cyclic amino acids, hydrogen bonding networks, constrained peptides and macro-cycles. In computational protein design, the JaLT algorithm can be used for dense sampling of loop conformations, for example of loops in active sites of enzymes in which a highly precise positioning of catalytic residues is required.

## 4 Methods

The three novelties of this paper are:

1. an algorithm to perform conformational change analysis of loops
2. a loop closure algorithm based on a Newton-Raphson scheme as in Ref. [35]
3. an inverse Jacobian analysis, which is the enabler of above two algorithms

The inverse Jacobian analysis is structured around *tripeptides*, which are clusters of three residues in which the *ϕ* and *ψ* dihedral angles are the free DoFs, as for example also used by Al-Bluwi et al. [2], and illustrated in Fig. 12. The three novelties are described in detail in the below subsections, preceded by a brief introduction to Screw theory, which forms the foudnation for the algorithms.

**Figure 12:**
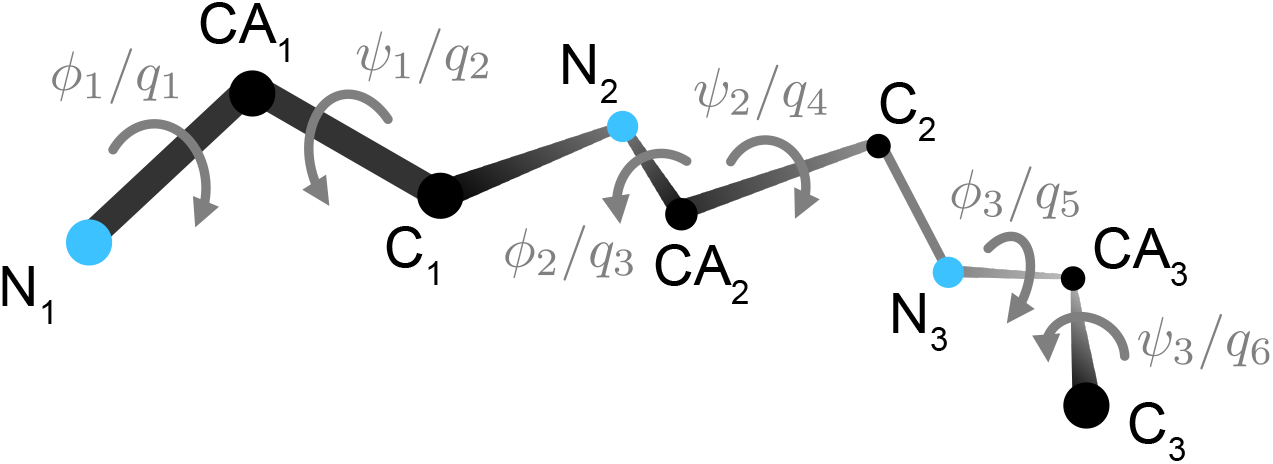
A tripeptide consists of three residues, where the *ϕ* and *ψ* angles are the free variables, i.e. the internal coordinates.

Throughout this section, scalars are indicated with an italic symbol, e.g. *d*, vectors are indicated with an arrow, for example 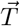, unit vectors are complemented with a hat, for example 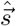, matrices are bold face, e.g. **J**, and lists of objects are contained in square brackets.

### Screw Theory

The algorithms introduced in this paper are built around a linearisation of the backbone kinematics, for which Screw theory is used. Screw theory is a form of linear algebra particularly well-suited to express the motion of bodies in Cartesian space. The motion of a body in Cartesian space has both an angular part and a translational part, and the linear algebra of such pairs of vectors is central to Screw theory. Huang et al. [24] specifically use the notion of twists and wrenches, where a twist 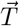 is a screw that consists of a Cartesian angular- and a translational motion vector, and a wrench 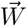 is a screw that consists of a Cartesian moment- and force vector:

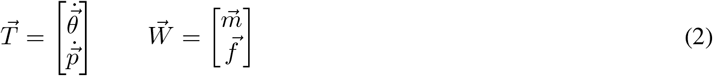

where 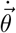 is the instantaneous rotational change vector of a body in Cartesian space, 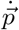 is the instantaneous linear displacement vector of a body in Cartesian space, and 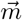 and 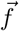 are respectively a moment and force vector acting on that body. It should be noted that 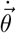 does not contain Euler angles, but is an instantaneous rotational change vector relative to the current orientation.

The last part of this method section introduces the Jacobian analysis of a tripeptide backbone that enables the mapping of a twist onto the torsion angles of a tripeptide:

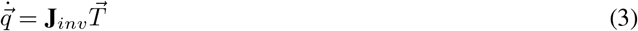

where 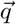 contains *ϕ_i_* and *ψ_i_* for *i* = 1․3 as shown in Fig. 12, and **J***_inv_* is the inverse Jacobian matrix that maps twist 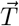 onto 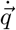. This linear transformation from Cartesian space to internal coordinate space is at the heart of the developed algorithms introduced below.

### Conformational change simulation algorithm

Before the details of the inverse Jacobian analysis are presented, a loop conformational change algorithm is presented. The latter algorithm perfectly demonstrates the advantages that an inverse Jacobian analysis brings to the field of protein analysis. As mentioned in the Introduction section, existing approaches to conformational change analysis often only use differences in torsion angles to guide sampling. The idea here is to combine that perspective on the difference between conformations with an expression of that same difference expressed in Cartesian space. The idea is that this maximizes the use of available information. A conformational change algorithm that includes an expression of the desired change in Cartesian space has two benefits:

1. the method can simultaneously solve expressions for the desired conformational change and expressions that represent the closure constraints, since both are expressed in a Cartesian vector space.
2. all Cartesian vectors can be expressed in a single Cartesian reference frame, which makes it possible to use linear algebra to weigh different vector that represent the same change. This gives a large degree of control over the sampling of the redundant DoFs in a protein loop.

The algorithm is introduced using pseudo-code in Algorithm 1 and starts with an Initialisation section.

#### Initialisation

Algorithm 1 starts with reading the user-defined options as discussed in the Results section. Next, the linear model of the loop is initialised, where the loop is divided into *N* segments and stored into a [*seg*] object. Each segment is a tripeptide, except for the final segment which may consist of one, two, or three residues, since not every loop contains an exact multiple of three residues. The N-atoms that connect subsequent segments are marked as *key atoms*, and their position and orientation form a non-redundant Cartesian representation of the loop conformation. The N-atom that follows the last segment must satisfy the closure constraints and its desired Cartesian change is therefore always zero (see also line 14-15 of Algorithm 1).

To ensure a one-to-one mapping of the error twist of a key atom onto torsion space, the minimum loop length is six residues. Six residues means two tripeptides and therefore one key atom that is not subjected to closure constraints. In other words, a six-residue loop has 12 dihedral DoFs, six of which are required to satisfy the loop constraints, which leaves six DoFs for unconstrained motion of the single key atom in Cartesian space. Beyond twelve residues, this algorithm accepts loops with any length.

#### Conformational change twists

The conformational difference between a provided start- and a target conformation can be expressed both in Cartesian space and in the internal coordinate space. The algorithm introduced in this paper considers both errors simultaneously. In each iteration the algorithm cycles through the different segments and calculates for that segment both the Cartesian error of the related key atom as well as the error between the respective torsion coordinates. In this paper all Cartesian positions and orientations are expressed in the reference frame connected to the first N-atom of the loop, which is the fixed basis for the loop. For each segment the desired change is a weighed combination of:

1. the difference in torsion angles between the current- and target conformation (line 11).
2. the difference in Cartesian coordinates of the key atom at the end of the segment (line 13).

Line 12 represents the transformation of the torsion angle difference onto Cartesian space, which is an intermediate step to project this difference onto the internal coordinates of other segments. The latter is required to define compensating moves to maintain loop closure. **J***_i,fw_* in line 12 is the forward Jacobian for the *i*th segment. The Jacobian matrices are updated at the start of each move (line 10), which will be discussed later. 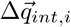 is calculated as the finite difference between the target values and the current values of the torsion angles that make up the *i*th segment,

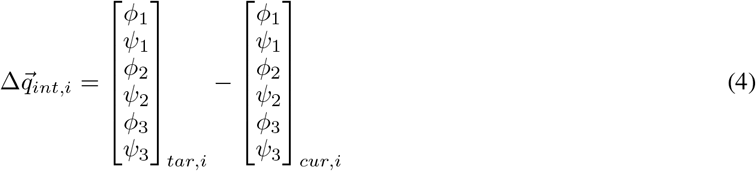

In the implemented function behind line 11 of Algorithm 1 the difference is mapped onto the range between −*π* and +*π* to handle the periodic nature of the internal coordinates. The notation 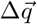 is used instead of 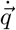 to indicate that finite torsion changes are being calculated in the iterative algorithm.

The Cartesian difference of the key atom at the end of each segment is calculated in parallel. The first step to generate a move for a key atom is to calculate its desired conformational change twist 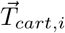. The twist 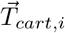 consists of an angular vector and a linear vector and its calculation is laid out in the `changeTwist` function in Algorithm 2. First the rotation matrix and translation vector, together referred to as an *RT* object, of the key atom *i* with respect to the global reference frame (the first N-atom of the loop) is calculated for both the current conformation and the target conformation (lines 2-3 of Algorithm 2). Then, the relative rotation and translation between these *RT* objects is expressed as *RT_rel_* (line 4). From *R_rel_* the delta ZYX-Euler angles are extracted (line 5), which are subsequently projected on Cartesian space using a `eulerZYXfromR` function (line 6). The latter represents the mapping

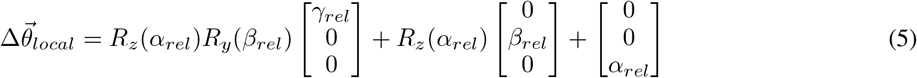

where *α_rel_*, *β_rel_*, and *γ_rel_* are the angles around respectively the *Z*, *Y*′ and *X*′′ Euler axes to represent the rotation matrix *R_rel_* matrix. It should be noted that the projection as expressed in Eq. (5) is a coarse approximation because Euler angles can be large, which makes the linear mapping potentially very inaccurate. However, this approximation becomes more and more accurate as the algorithm progresses and the rotation matrix in *R_rel_* approaches the identity matrix, at which point the three *ZY X* Euler axes align with the three Cartesian unit vectors. This conversion to orthogonal axes is beneficial for algorithm conversion and the reason for choosing the *ZY X* convention.

**Algorithm 1:**
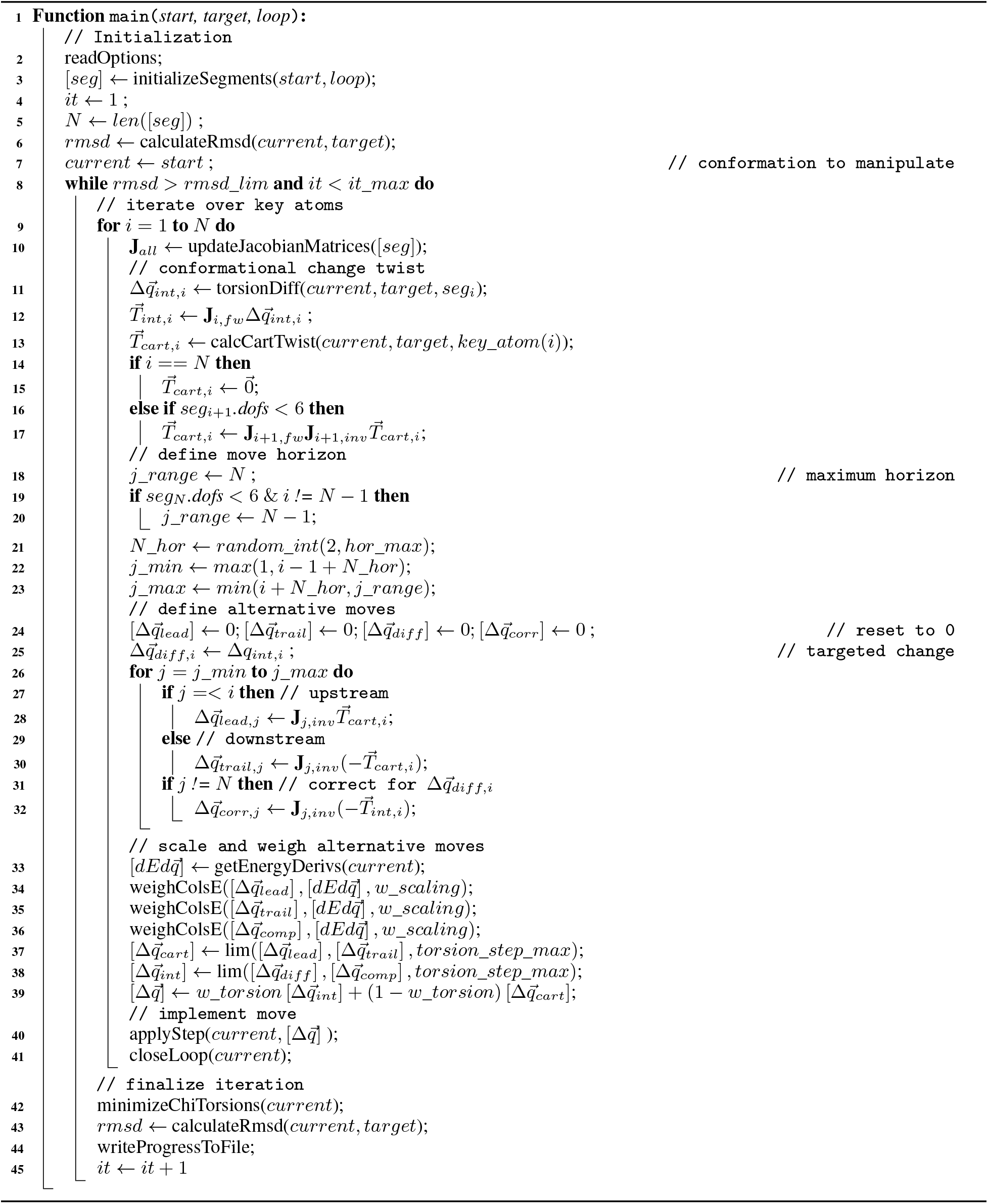
Conformational change analysis

**Algorithm 2:**
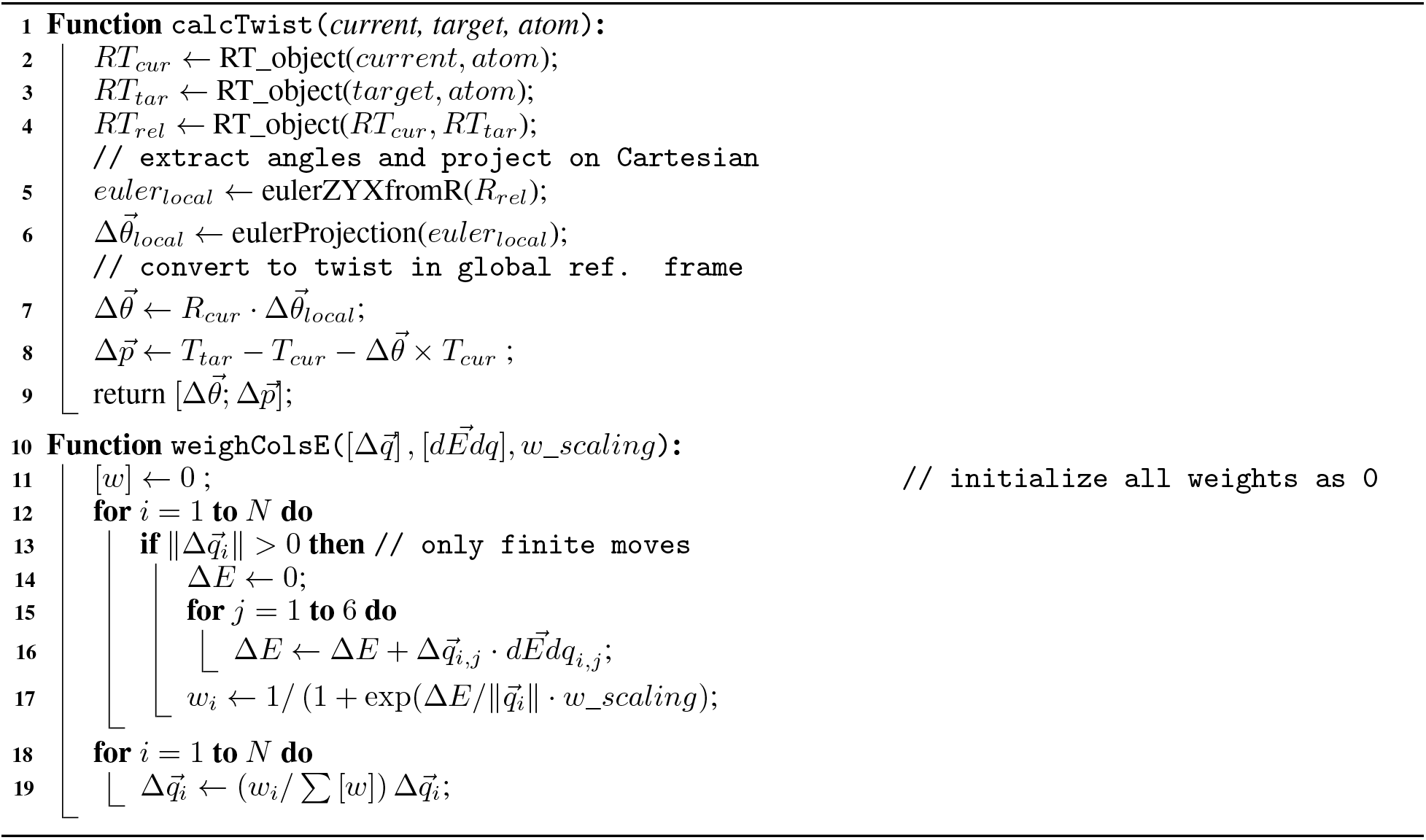
Support functions

The resulting vector 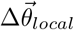 is expressed in the local reference frame, and thus needs to be transformed into the global reference frame (line 7 of Algorithm 2). The same holds for the relative translation vector between the *RT* objects, in which case the induced rotation needs to be accounted for (line 8 of 2). The vector with projected Euler angles and the translation vector together form the instantaneous, finite error twist (line 9 of 2), which is expressed as 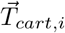 in line 13 of Algorithm 1.

There is one final consideration in the calculation of the change twist. In order to maintain loop closure, the segments that trail a key atom need to realize the opposite of the defined change twist. However, in case a loop cannot be divided in an exact number of tripeptides, the final segment will be a dipeptide or a monopeptide and not span six-dimensional space. In that case the final segment may not be able to generate the required compensatory motion to satisfy the closure constraints. To prevent this, 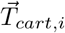 is mapped onto the Cartesian space of its trailing segment (*i* + 1) in case it has < 6 DoFs. For that 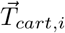 is first mapped onto the linearised torsion space using the inverse Jacobian, and subsequently mapped back onto Cartesian space using the forward Jacobian (line 16-17 of Algorithm 1). The key atom at the end of the last segment is a loop connection atom and thus remains fixed throughout the simulation. Therefore, 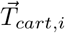 for the final segment is always the zero twist (lines 14-15).

#### Move horizon

For a given key atom the segments that are leading that key atom are upstream in the kinematic chain and changes in their torsion angles contribute to the motion of the considered key atom. On the other hand, changes in the torsion angles of tripeptides that trail the key atom do not induce motion of the key atom, but are instead used to maintain kinematic closure of the loop. While in theory all torsion angles of a loop can be involved in a key atom move, it is both necessary and beneficial to be able to control the *horizon* of a move. The move horizon is defined here as the maximum number of segments on either side of a key atom that are allowed to contribute to the move.

Firstly, it is necessary to control this horizon because final segments that are a monopeptide or dipeptide should generally be excluded from a move since they span a space with less than six DoFs. If the tripeptides on either side of a key atom span six-dimensional space, then the final segment is always excluded if it does not span six-dimensional space (lines 18-20 of Algorithm 1).

Secondly, it is beneficial to limit the horizon of a move to prevent excessive interference between moves, e.g. a move on one end of the loop reverts progress achieved on a key atom at the other end of the kinematic chain. For this reason the user option `hor_max` was introduced, which defines the maximum number of segments on either side of a key atom that are involved in a move. The actual number of segments on either side of a key atom is for each move randomly picked as a number between 2 and `hor_max` (line 21). This is the only randomness in the algorithm and makes that each run generates a different conformational change trajectory. The resulting value *N*_*hor* subsequently defines the lower- and upper move horizon (lines 22-23).

#### Alternative moves to realize change twist

With the conformational change twists calculated and the move horizon defined, the next step is to calculate the set of torsion change vectors that represent the conformational change twists in internal coordinate space. This is done differently for 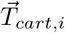 and 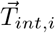.

In case of 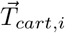, the segments within the horizon that are leading the key atom are responsible for generating the change twist. Their torsion change vectors are calculated by mapping 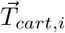 onto the torsion space using the inverse Jacobian matrix (line 28 of Algorithm 1). The matrix **J***_j,inv_* refers to the matrix resulting from Eq. (13) for tripeptide segment *j*. The segments within the horizon that are trailing the key atom are responsible for generating the opposite of the change twist, so that the net motion of the final atom in the loop is zero. As such, their torsion change vectors are calculated by mapping 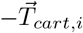 onto the torsion space using the linearised kinematic mapping (line 30).

The twist 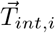 represents a desired change of the torsion angles 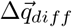 in segment *i*, as calculated using Eq. (4). 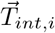 can therefore be considered as the Cartesian representation of the desired torsion change in segment *i*. To maintain closure, this change needs to be compensated by one or more other segments within the horizon, but it does not matter whether they are leading or trailing the considered segment. As such, for each segment within the horizon except for segment *i*, the compensating torsion vector 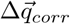 is calculated by mapping 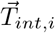 onto the respective torsion space (lines 31-32), where the final segment (which may have less than 6 DoFs) is always excluded for simplicity.

The resulting vectors-of-vectors 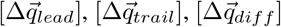, and 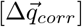 each contain *N* vectors, of which all vectors outside the horizon are zero vectors. Moreover, the non-zero entries in 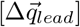 and 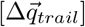 do not overlap, and neither do the non-zero entries in 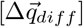, and 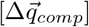.

#### Weighted and limited combination of moves

Because each alternative torsion change vector represents 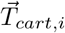 or 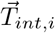 in local internal coordinates, weighing needs to be applied so that the sum of all the weights is equal to unity for both 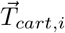 or 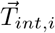 (line 34-36 of Algorithm 1). Weighing is performed based on the energy gradient of each alternative move.

For each alternative torsion change vector, i.e. the entry has a non-zero norm (line 13 of Algorithm 2), the energy gradient of the net expected energy change is calculated based on the linearised system (line 14-16 of Algorithm 2). In the Rosetta software suite the energy gradient with respect to a torsion angle change is calculated using the method outlined by Abe et al. [1]. The resulting Δ*E* is normalized with respect to the norm of the torsion change vector, which is the effective energy gradient of the torsion change vector in question. Because the energy function is highly non-linear, a customisable sigmoid function is used to normalize the effective energy gradient (line 17 of Algoirhtm 2). The chosen sigmoid function is a negated logistic function, so that negative energy gradients get a higher weighing, of which the spread can be controlled with the user option `w_scaling`.

Because this algorithm calculates moves based on a linear representation of the kinematics, which is only accurate in the direct proximity of the instantaneous conformation, the step size needs to be limited. For this reason Jacobian matrices are updated before each key atom move (line 10 of Algorithm 1), but as an additional safe-guard the calculated torsional change vectors move are also normalized based on a maximum allowed change per move; Δ*q_max_* (line 37-38 of Algorithm 1). After limiting both the Cartesian-defined move (based on 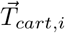) and the internal coordinates-based move (defined by 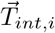), they are linearly combined based on the user-defined, relative weight factor `w_torsion`.

#### Iteration finalisation

The final step of each iteration in Algorithm 1 is the implementation of the weighted and limited torsion change vectors that together form the move for a segment and its related key atom. If the system would be perfectly linear, then the move would maintain perfect loop closure thanks to the trailing move changes (in case of Cartesian change vector formulation) and the compensating move changes (in case of internal coordinate change vector formulation), which realize the opposite twist. However, protein kinematics are far from linear, and therefore kinematic closure is easily disrupted. For this reason a second iterative algorithm is introduced, which enforces kinematic closure and thereby supports the conformational change algorithm. That algorithm is introduced in the next section.

Once the conformational change algorithm has iterated over all segments with related key atoms, it ends with a call to a side-chain energy minimisation algorithm. The minimisation algorithm is an Armijo line search where the variables are all *χ*-angles of all residues in the loop, as well as those of all residues within a range of 10 Å. Side-chain minimisation allows a meaningful comparison of the sampled backbone conformations based on their Rosetta energy score, which is essential to analyse the kinetics of loop conformation changes.

### Loop closure algorithm

The novel iterative, Jacobian-based kinematic closure algorithm is described in Algorithm 3. It builds on the concept of a series of tripeptide segments, which was captured by the [*seg*] object, and also reuses the `calcTwist` function that was introduced in Algorithm 2. In this case, however, there is only one atom of interest and that is the final C-atom of the loop. Since only *ϕ* and *ψ* torsion angles are free, the link between the last C-atom in the loop and the subsequent N-atom is not changed by the loop closure algorithm, because it concerns an *ω* angle.

Therefore, a successful kinematic closure means that the relative position and orientation of the final C-atom with respect to the first atom of the loop is the same as in the *start* conformation. The twist that represents this loop closure error is indicated by 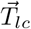. The kinematic closure algorithm iterates until the norm of both the rotation vector and the translation vector of 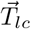 have been brought below a pre-defined threshold, or a maximum number of iterations is reached. The thresholds are set at *ϵ*_*lin* = 0.1 Å, and *ϵ*_*rot* = 0.087 rad (5 degrees). The maximum number of iterations is set to three iteration to ensure that closure solutions are very close to the current conformation. The latter is necessary to ensure that the closure algorithm does not interfere with the overall progress in conformational change that is managed by Algorithm 1.

Each iteration step starts also here with an update of the Jacobian matrices (line 5). Next, for each segment the loop closure twist is mapped onto the linearised torsion space (lines 7-8). Similar to Algorithm 1, each vector forms an alternative solution to the loop closure, so a weighing is required. In this case the goal is to achieve closure with minimal disturbance to the loop conformation, whereas in Algorithm 1 the goal was energy minimisation. To penalise disturbance to the loop conformation, the different torsional change vectors are weighed based on the inverse of their squared norm (line 25).

In case the last segment is not a tripeptide (and thus does not span six-dimensional space), a correction is required. In that case the mapping of line 8 is not one-to-one, which introduces an error. This error can be expressed in Cartesian space by applying the forward Jacobian on the torsion change values associated to the assisting residues of that segment (line 11). As an example, in case the final segment is a dipeptide (i.e. four DoFs), then the **J**_*fw,cons*_-matrix will be a 6 × 6 matrix where only the last two columns are occupied. Once expressed in Cartesian space, this correction twist 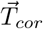 can be mapped onto the torsion space of all tripeptide segments (lines 13-14), weighed (line 15), except for the final segment, which is corrected for and for which therefore a negative mapping is applied (line 16). Finally, the correction vectors are added to the earlier defined torsion change vectors (line 17). Each iteration ends by applying the calculated torsion changes to the conformation, followed by an update of the loop closure twist to determine whether the algorithm has converted to within the specified bounds.

**Algorithm 3:**
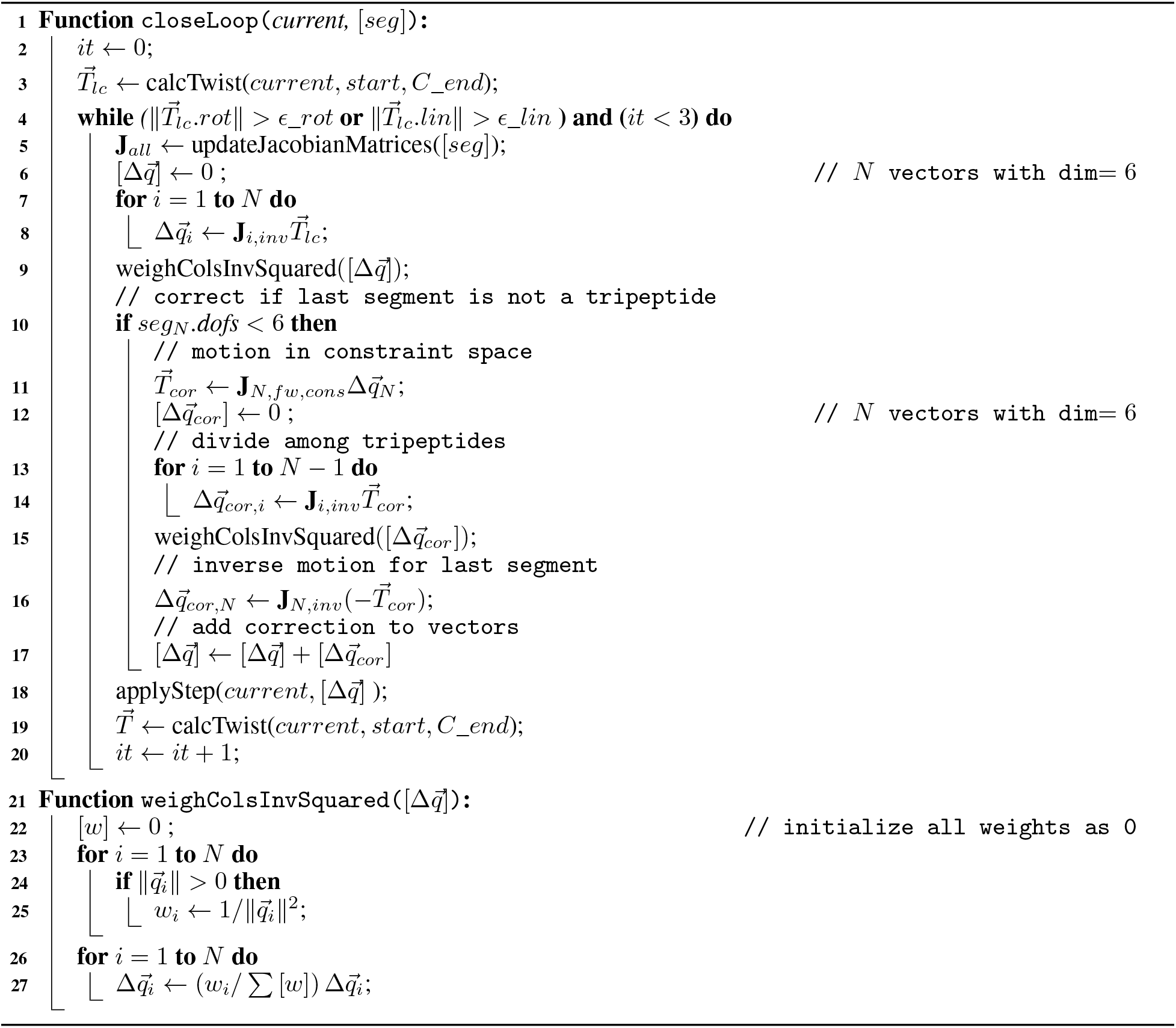
Inverse Jacobian loop closure

### Inverse Jacobian analysis of tripeptides

It was mentioned earlier that both the conformational change algorithm and the loop closure algorithm updates the Jacobian matrices at the start of each move (line 10 of Algorithm 1). This section describes how that is done. To perform the inverse Jacobian analysis of a tripeptide, this paper applies the *generalized Jacobian analysis* method that was developed by Huang et al. [24]. They based their method on Screw theory as briefly introduced earlier in this Method section.

Both twists and wrenches are six-dimensional objects whose bases can be freely defined in dual Cartesian space. In the generalized Jacobian analysis these bases are defined such that each wrench is orthogonal to all twists except for one (see Huang et al. [24] for more details). In this paper the basis twists and basis wrenches are described for a tripeptide with the dihedral angles *ϕ* and *ψ* as the free variables so that each tripeptide forms a six DoF backbone segment.

The basis for the twist space is defined by the rotation axes of the three pairs of *ϕ* and *ψ* torsion axes,

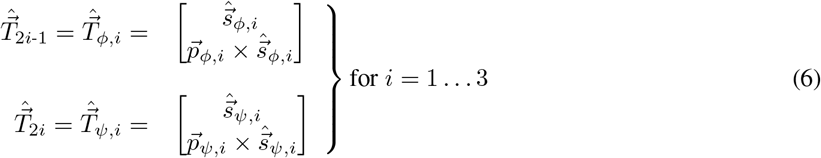

in which 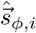 and 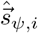 are the unit vectors along the *ϕ* and *ψ* torsion axes of the *i*th residue of the tripeptide, and 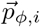 and 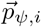 are position vectors from the origin of the reference frame to a point on the two respective axes. See Fig. 13 for examples of the vectors. It needs pointing out that the analysis presented here does not depend on the chosen reference frame, as long as all vectors are expressed consistently in the same reference frame.

**Figure 13:**
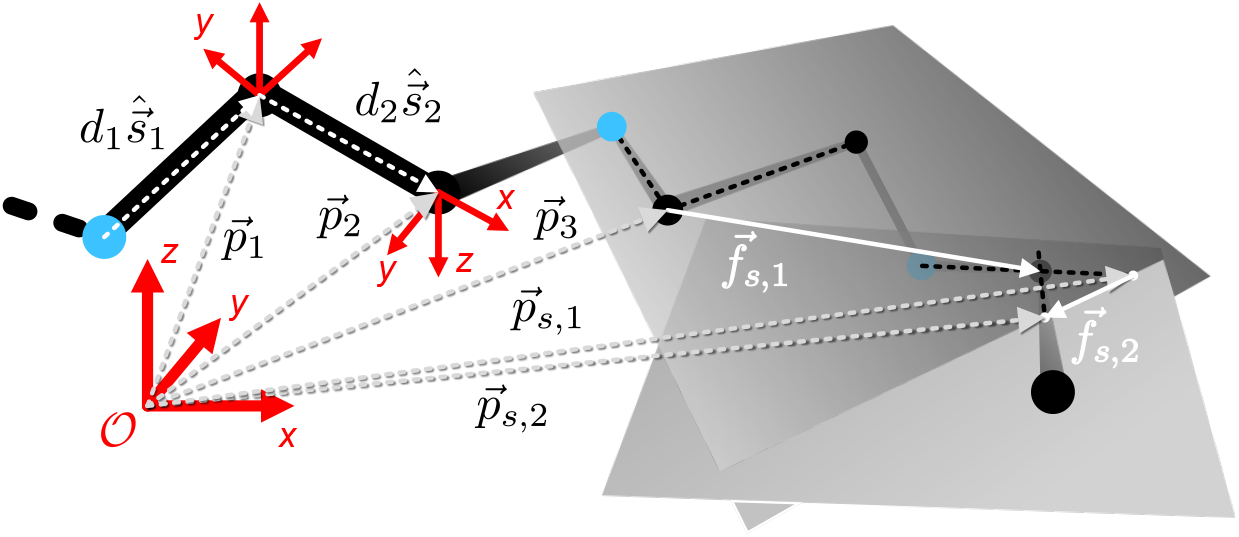
To perform the Jacobian analysis of a tripeptide, the vectors connecting atoms are described as the combination of a unit vector 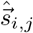 and a magnitude *d_i,j_*, which is the bond length. Various position vectors are indicated with 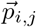.

In the nominal case all six basis twists are independent. A discussion on singular configurations is included later in this section. When all six basis twists are independent, any arbitrary twist can be expressed as a linear combination of the six basis twists,

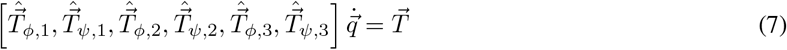

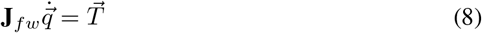

where **J***_fw_* is the forward Jacobian matrix constructed using the six basis twists, and

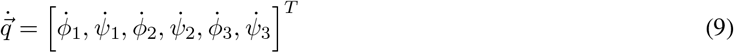

where 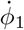 to 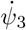 are the magnitudes of the tripeptide’s six torsion gradients in the internal coordinate space.

The generalized Jacobian analysis relies on the definition of a basis for the wrench space such that each wrench is dual to (does work on) exactly one twist, and thus is orthogonal to the remaining five twists. If unit wrench 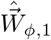 is orthogonal to all unit twists except 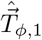, then left-multiplication of Eq. (7) with the transpose of the wrench 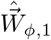 gives

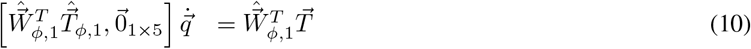

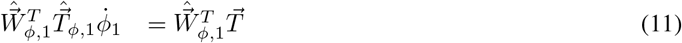

The transposed wrench multiplied by the twist at the left side of Eq. (11) is a scalar, so therefore Eq. (11) can be rewritten as

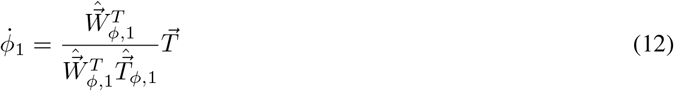

which is the inverse Jacobian analysis for the torsion angle *ϕ*_1_. Repeating Eqs. (11)–(12) for the other five torsions and combining the results gives

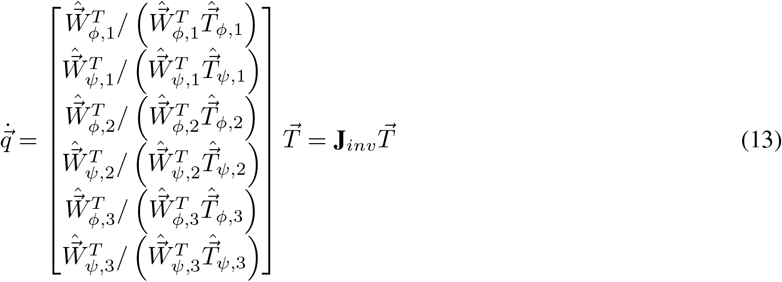

where **J***_inv_* is the inverse Jacobian matrix for a tripeptide. The difficulty lies in the definition of the wrenches in Eq. (13), which is done in the next section.

#### Definition of dual wrenches

The twists that are required to perform the analysis in Eq. (13) were defined in Eq. (6). This subsection derives expressions for the wrenches based on geometric insight, which is done in two steps.

The first step is the definition of three sets of two wrenches, where the space spanned by each set of two wrenches is dual to a set of two twists, and therefore is orthogonal to the other four twists. To allow for more specific description, the two wrenches that are dual to 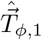 and 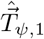 are described as an example. A first wrench is identified as the force that acts along the line that connects points CA_2_ and CA_3_, illustrated with the support vector 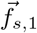 in Fig. 13.

The second wrench in the set acts along the intersection line of the two planes defined by respectively the *ϕ*_2_ and *ψ*_2_ torsion axes and the *ϕ*_3_ and *ψ*_3_ torsion axes. For the example of the twists 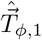 and 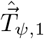, this intersection line is expressed using vector 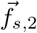 as shown in Fig. 13. To define this vector, two supporting points are used, namely 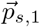, the intersection of the torsion-axis belonging to *ϕ*_3_ with the plane defined by the *ϕ*_2_ and *ψ*_2_ torsion axes, and 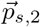, which is located at the intersection point of *ψ*_3_ with the plane defined by *ϕ*_2_ and *ψ*_2_ torsion axes. The line connecting these points is labelled 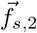, and is shown in Fig. 13 together with examples of all other vectors described above.

A mathematical expression for the location of 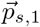 and 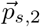 can be obtained using linear algebra. The analysis presented here is based on a system where the reference frame connected to each atom has its X-axis aligned with the incoming bond, it’s Z-axis as the cross product of the before-last bond vector with the incoming bond vector, and Y-axis complementing a right-handed system. Figure 13 shows the reference frames connected to CA_1_ and C_1_ as examples. The analysis relies on the availability of a rotation matrix 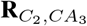 and a translation vector 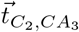 which together transform a vector from the reference frame connected to CA_3_ in the reference frame connected to atom C_2_ as

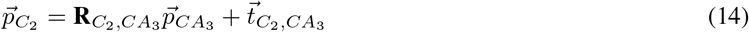

where 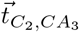 is a vector expressed in the reference frame connected to the atom CA_3_ and 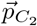 is a vector expressed in the reference frame connected to the atom C_2_. If these objects are expressed numerically, then 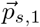 is located on the 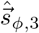-axis at a distance

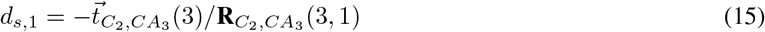

where the numbers between brackets indicate a row and column index. Equivalently, 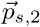 is located on the 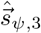-axis at a distance

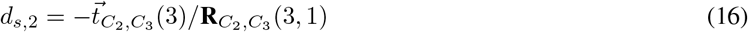

where 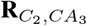 and 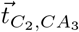 are the rotation matrix and translation vector which together transform a vector from the reference frame connected to C_3_ in the reference frame connected to atom C_2_.

Now, the two unit wrenches that span a space that is dual to the space spanned by twists 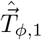 and 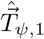 can be expressed as

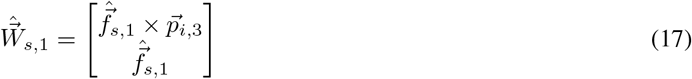

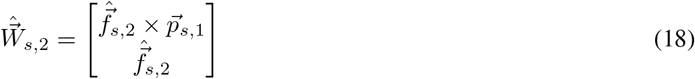

where 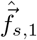 and 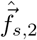 are the normalized versions of 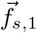 and 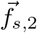.

The second step is to find two normalized combinations of 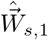 and 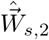,

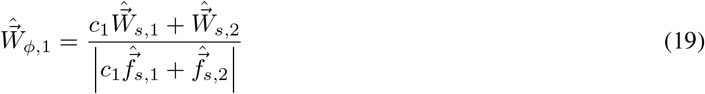

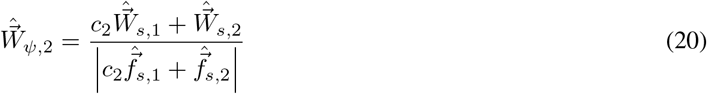

so that 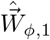 is orthogonal to 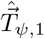 and 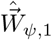 is orthogonal to 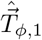, i.e.

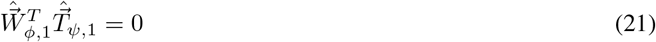

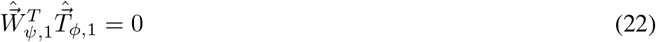

Combining Eqs. (19) and (21), and Eqs. (20) and (22) results in

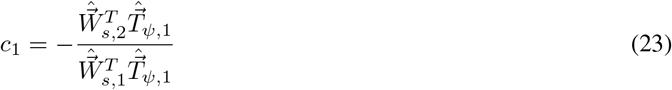

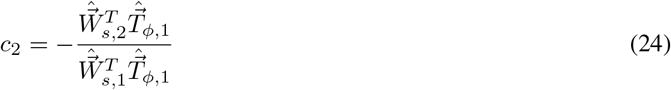

Notice that Eqs. 19 and 20 are normalized based on the force vector (the second of the wrench’s vectors), which is independent of the coordinate frame in which the twists and wrenches are expressed. A similar analyses is performed for the other pairs of twists and wrenches.

#### Singularity detection

The main advantages of performing an inverse Jacobian analysis using linear algebra is that it allows detection and handling of singularities. Singularities are the main reason that an inverse Jacobian matrix cannot be obtained by taking the numeric inverse of a forward Jacobian matrix. Singularities arise when a Jacobian is rank-deficient, which can be caused by various reasons:

- *Coplanar sets of twists.* In case any of the planes described by a set of *ϕ* and *ψ* axes is parallel to one of the other planes, then an intersection point cannot be defined. This can be detected by evaluating whether the denominators of Eq. (15) and (16) are close to zero. If this is detected, the algorithm handles this by replacing the near-zero value with a small finite value, essentially creating an artificial intersection point a long distance away.
- *Dependent twists.* If two (or more) of the six torsion axes of a tripeptide align, then the tripeptide no longer spans the full Cartesian space. This results in dependent rows in Eq. (13) and means that there are Cartesian vectors that are mapped on the null space of the tripeptide’s torsion space. As such, the mapping is no longer one-to-one.

Since there is nothing that prevents a protein from transitioning through kinematic singularities, and protein motions involve a large number of motion steps, it is essential that a computational simulation method can handle singularities. As opposed to numeric inversion of a forward Jacobian matrix, which fails when the Jacobian is rank-deficient, an inverse Jacobian analysis based on linear algebra can be made robust against such singularities and therefore acts as an enabler for kinematic motion analysis of protein loops.

#### Constrained tripeptides

Only in special cases can a loop segment be divided in an exact number of tripeptides. If that is not the case, than the final segment will be a monopeptide (2 DoFs) or dipeptide (4 DoFs), and thus not span six-dimensional space. To represent those constraints, this final segment is complemented with two (in case of a monopeptide) or one (in case of a dipeptide) *assisting* residues, borrowed from the preceding segment. This allows six twists and wrenches to be defined, but those associated to the assisting residues now represent the motion space that is inaccessible to the segment in its instantaneous configuration. This concept and the related numbering of internal coordinates is shown in Fig. 14.

**Figure 14:**
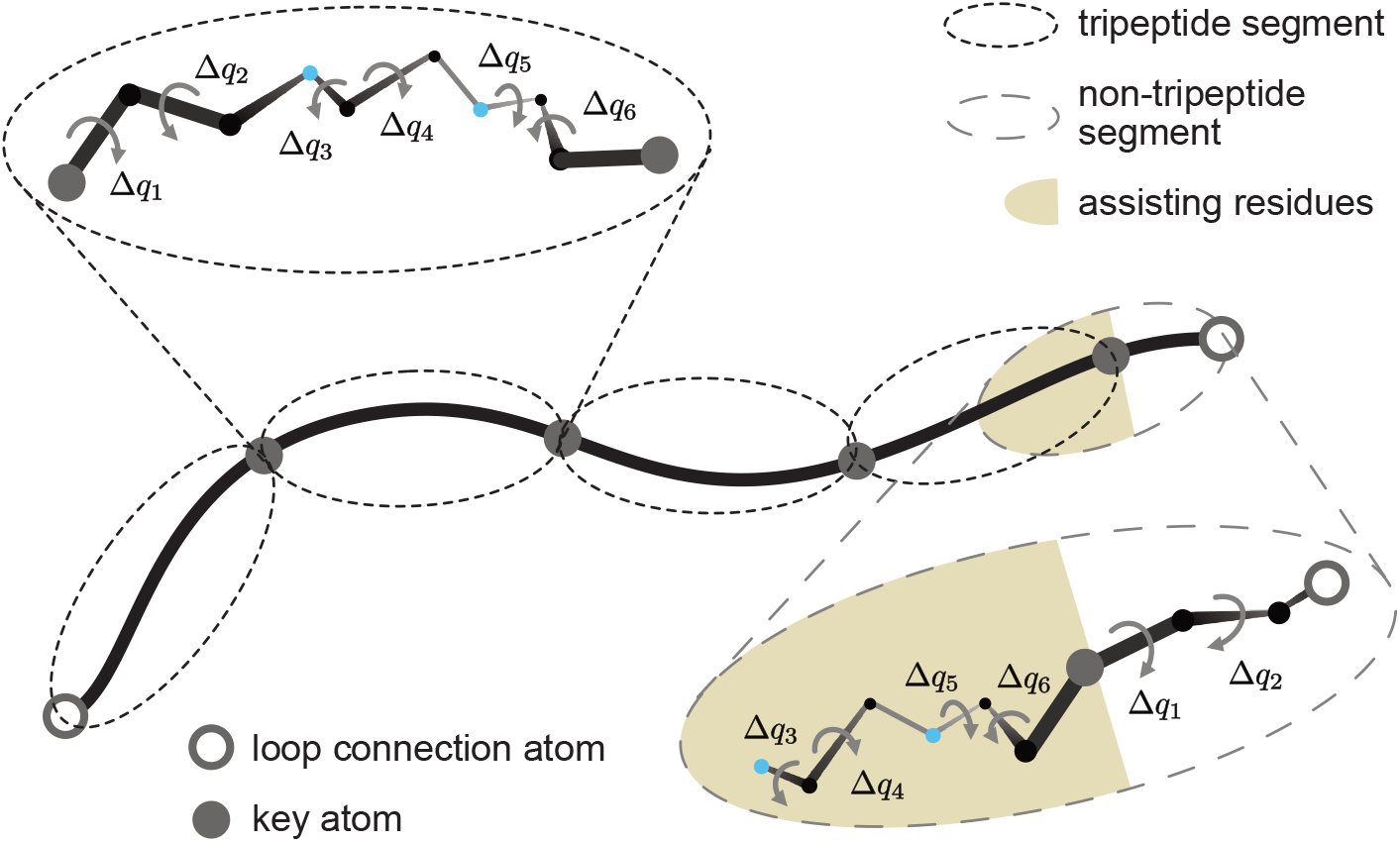
For the presented analyses a loop is divided in segments which by default are tripeptides (i.e. 6 DoFs), unless it is the final segment. In the example shown the final segment consists of a single peptide (i.e. 2 DoFs). In such a case, the two preceding peptides are added to represent the constrained space of that segment, which spans the remaining 4 DoFs for this example.

### PDB preparation

The simulations presented in this paper were run on the MET20 loop of DHFR. The PDB with identifier 1ra2 was used as starting conformation for the open state and the PDB with identifier 1rx7 was used as the target conformation for the closed target state. The PDBs were subsequently:

1. reduced to a single chain
2. relaxed in Rosetta using the command `main/source/bin/relax.macosclangrelease-relax:constrain_relax_to_start_coords` -`relax:coord_constrain_sidechains-relax:ramp_constraints false` -`out:file:renumber_pdb true`.
3. cleaned by removing all ligands. For this all HET lines were removed from the PDB
4. aligned to each other so that PDBs could be directly compared in PyMOL

The loop spans residues 9 (Ala) to 24 (Leu) in the resulting PDBs.

**Table 3:** Supplemental files provided with this research.

| Name | File | Description |
| --- | --- | --- |
| Supplemental Data 1 | verification_data.zip | contains the simulations results that were used to analyse the performance and characteristics of the algorithm as described in the section 'Algorithm performance' |
| Supplemental Data 2 | met20_simulations.zip | contains the data from all simulations that got to within 1 Å CA RMSD of the target conformation within 500 iterations steps. This amounts to 244 of the 2000 started simulation for the wild-type 1ra2, and 272 of the 2000 started simulation for the I14G-mutated variant. For each simulation per iteration step the Rosetta score (REU), the values for the CA RMSD to both 1ra2 and 1rx7 (Å), the torsion angle RMSD values to both 1ra2 and 1rx7 (deg), and the torsion angle for all $\phi$ and $\psi$ angles of residues 9-24 (deg) can be found in the data files. |
| Supplemental Data 3 | example_trajectories.zip | contains the pre-processed 1ra2 and 1rx7 PDB input files as well as PDBs for the intermediate states of the example trajectories presented in Figs. 9 and 11. Intermediate states were generated at intervals of 10 iterations. |
| Example Input | example_input.zip | contains a text file with an example command to run the *loop_transition* application, together with examples of required files as input. The first required file is a .txt file with, on separate lines, the path to respectively the start conformation, the target conformation, a Rosetta loop definition file for the start conformation, and (optional) the corresponding loop definition file for the target conformation (in case residue number deviates). As such, the files referred to in the .txt file are also input files, and examples are added. N.B. the output command (following the '-o command) does not need an extension. |

## Supporting information

Example Input

## Acknowledgement

The authors would like to thank the Rosetta Community for funding this research and its members for the many insightful discussions throughout the development of the ideas that are presented in this publication.

## Code and data availability

The code will be distributed with in Rosetta macromolecular modelling package. In anticipation of this, an example input can be found in the supplemental Example Input. For information about how to obtain access to the Rosetta code, please visit https://www.rosettacommons.org/software. As part of this manuscript, the following files are included as supplementary information:

